# Brain state locks silent memories in a trained fly connectome

**DOI:** 10.64898/2026.10.24.761586

**Authors:** Han Wang, Joe Yeong Poh Sheng, Yunhua Zhao, Yuhui Zhao, Zizhong Yu, Xiaoyu Liu, Ruoyun He, Huikuan Chen, Ting Liu, Ye Tian, Weibang Chen, Yao Cai, Yue Xu, Hongyu Ning, Xuanhe Hou, Lingpin Pang, Danxian Jiang, Huafeng Liu, Junfeng Hao, Chunjie Tian

**Author notes:** Correspondence (Huafeng Liu). Correspondence (Junfeng Hao). Correspondence (Chunjie Tian). These authors contributed equally: Han Wang and Joe Yeong Poh Sheng.

## Abstract

Complete connectomes now exist for insect nervous systems, but whether a fixed wiring diagram can learn, and what its memories consist of, is unknown. Here, we trained a spiking model of the whole male *Drosophila* central nervous system, 167,106 neurons with fixed wiring, to plan moves and to remember a visual cue. Spikes write the cue into neuronal adaptation, which holds it silently until the next input reads it out. Cells that hold the trace are enriched for transcripts of a calcium-activated potassium channel in fly cell atlases. The learned readout recalls the trace only in the brain state of learning: the same trace evokes a different response at another delay, and a disease-like loss of potassium current rewrites the trace and changes how it is read. Learning in the new state is spared. Memory failure can arise from a change of state rather than a loss of storage.

**HIGHLIGHTS:**

- A fixed whole-fly connectome learns to plan and to remember a visual cue
- The cue is held silently in neuronal adaptation and read out by the next input
- Recall works only in the brain state of learning, at writing and at reading
- A disease-like lesion rewrites the trace, while learning in the new state is spared

**IN BRIEF:** Wang et al. train a spiking model of the whole male fly nervous system to plan and to remember. It holds a visual cue silently in neuronal adaptation and recalls it only in the brain state of learning, so a disease-like lesion removes recall while new learning is spared.

## INTRODUCTION

Complete connectomes now map every synapse of an insect’s brain and nerve cord, ^1–3^ and their cell types can be matched across animals and between the sexes. ^4,5^ Models built on them predict sensorimotor pathways and neural activity, either without training or with training confined to one sensory pathway. ^6–8^ In a physics model of the fly’s body, trained controllers walk and fly, but they are not built from the fly’s wiring. ^9^ The connectome itself is often described as a static map without dynamics, state or plasticity. ^10^ Whether a fixed wiring diagram can learn demanding behavior, and what its memories would be made of, is unknown.

How a brain holds information across a delay is still disputed. Persistent firing carries remembered items in recordings from monkeys and humans, ^11,12^ in areas from prefrontal to visual cortex, ^13^ whereas activity-silent states ^14^ can be read out by an impulse. ^15^ Whether silent states hold working memories, serve only passive storage or reflect retrieval from episodic memory is debated. ^16–18^ Such states are almost always attributed to synapses, through short-term facilitation or fast Hebbian plasticity, ^19–22^ while intrinsic states such as spike-frequency adaptation have been studied mainly as aids to computation. ^23^ In the fly, visual memories that last several seconds depend on signaling in ring neurons of the central complex. ^24,25^ Which substrate learning recruits on real wiring is open.

Where memories live, and why learned codes hold in some states but not in others, is equally unresolved. Engrams of a single memory are spread across many brain regions, ^26,27^ and the excitability of a neuron at learning biases whether it joins one. ^28,29^ Population codes can stay stable across a delay while responses to an impulse change with time; ^30,31^ stable and dynamic codes coexist in prefrontal cortex, ^32^ and in human visual cortex the code for a remembered location turns toward the coming eye movement. ^33^ Retrieval can require the brain state present at encoding. ^34,35^ Changes of excitability alone can break a decoder trained in one state, whereas a decoder trained across states keeps working. ^36^ Which part of a memory a change of state alters, the stored trace or its reading, has not been tested in a complete nervous system.

In Alzheimer’s disease, early memory loss can be a failure of access rather than of storage. In mouse models of early disease, activating engram cells recalls memories that natural cues no longer retrieve, ^37^ and activating parvalbumin interneurons restores retrieval. ^38^ Neuronal hyperexcitability is an early feature of the disease, ^39–42^ but lowering it has had mixed effects on cognition in patients. ^43–45^ In flies, amyloid-β42 (Aβ42) degrades the A-type potassium channel Kv4, ^46^ and the loss of Kv4 from mushroom-body neurons underlies part of the courtship memory deficit that Aβ42 causes. ^47^ Why a lesion that spares new learning removes existing memories is not understood. Each of these questions could be asked of a connectome that learns, in which every state variable of every cell can be recorded and moved between brain states.

Here, we trained a digital twin of the fly’s nervous system: the complete wiring of one male animal, simulated cell by cell. It was trained on tasks rather than fitted to recorded activity, checked against fly physiology and replayed bit for bit, with every experiment declared before it ran. The model learns to plan moves and to remember a visual cue. Spikes write the cue into neuronal adaptation, which holds it silently until the next input reads it out. The trace is spread across the nervous system, and no ensemble of cells is more necessary than matched sets. The cells that hold it are enriched for transcripts of *SK*, a calcium-activated potassium channel, in fly cell atlases. The learned readout uses the trace only at the delays it learned, because the same trace drives a different response at each delay. The wiring keeps the readout reachable under heavy cell loss, and training across excitability states widens access, as decoder theory predicts. A disease-like loss of potassium current after learning rewrites the trace and changes how it is read, so recall returns only in the state of learning, while learning under the loss is spared.

## RESULTS

### A whole-CNS connectome learns to plan and to remember

We built a spiking model of the complete central nervous system (CNS) of a male fly from its connectome: ^2^ 167,106 leaky integrate-and-fire neurons with spike-frequency adaptation, joined by 25.6 million fixed connections (Figures 1A and 1B). Training could change 35,310 gains and thresholds, set per cell type, and the 2,052 weights of a linear readout, but never the wiring (Figure 1E). The model sees each task through 3,200 visual input cells, and 3,555 of its 4,096 readout cells (87%) receive a synapse directly from them (Figure 1E). Training reached the whole CNS: it changed the output gains of nearly every cell, in every division (Figures S1A–S1C).

**Figure 1.**
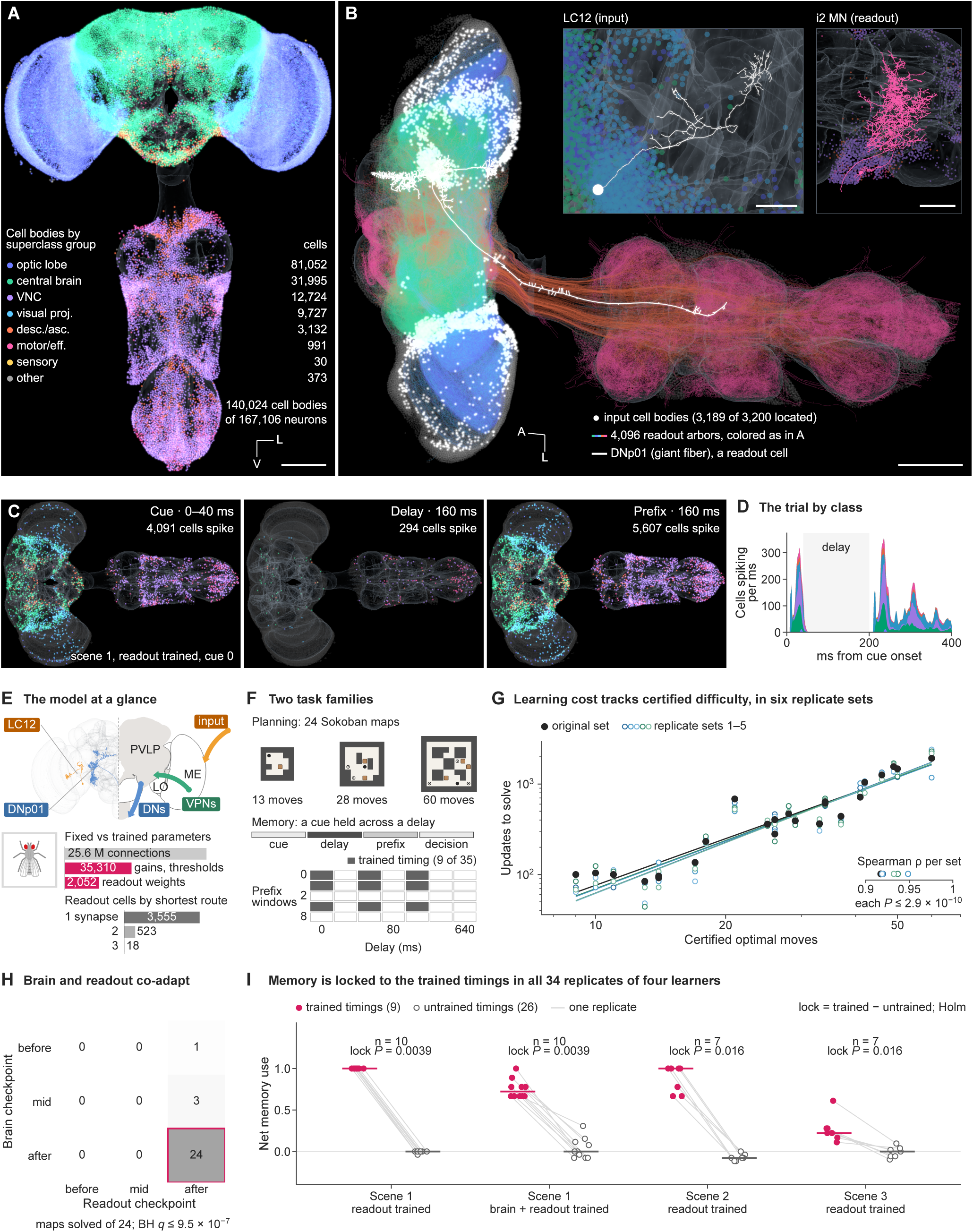
A whole-CNS connectome learns to plan and to remember. (A) The unfolded CNS of the model: every located cell body, colored by superclass group. Scale bar, 100 μm. (B) The interface: the visual input cells (white), the arbors of the 4,096 readout cells and the giant fiber DNp01; insets, an input cell (LC12) and a readout cell (i2 MN). Scale bars, 100 μm; insets, 20 μm (LC12) and 50 μm (i2 MN). (C and D) One memory trial (scene 1, readout trained, cue 0): the cells that spike in the cue, the delay and the input (C), and the cells spiking per ms by group (D). (E) The model at a glance: input and readout pathways, fixed and trained parameters, and the readout cells by their shortest route from the input. (F) The tasks: three of the 24 planning maps with their optimal moves, and the delayed-cue protocol with its 9 trained of 35 timings. (G) Learning cost against certified difficulty, 24 maps in 6 replicate sets; lines, least-squares fits on log axes; inset, Spearman ρ per set with permutation tests. (H) Maps solved by each pairing of brain and readout checkpoints; McNemar tests, BH-adjusted. (I) Net memory use at trained (filled) and untrained (open) timings in 34 replicates of four learners (10, 10, 7 and 7), joined by replicate; bars, medians; exact sign-flip tests of the lock (trained minus untrained use), Holm-adjusted. See also Figure S1 and Video S1.

We trained the model on two tasks (Figure 1F). To plan, it pushes boxes onto targets in a ladder of 24 maps of rising difficulty, each with a certified optimal solution. To remember, it is shown which of two targets is cued, holds the cue through a blank delay of up to 640 ms and then, from the board without the target, pushes the box toward the cued one. During a memory trial, cells throughout the CNS spike in the cue and the input, and few spike in the delay (Figures 1C and 1D).

The model learns both tasks reproducibly. Learning cost tracked the certified difficulty of the maps in all 6 replicate sets (Spearman ρ = 0.917 to 0.949; Figure 1G; Figures S1D and S1E). The final brain with the final readout solves all 24 maps, whereas no other pairing of brain and readout checkpoints solves more than 3 (Figure 1H), so brain and readout learned together; Video S1 shows the hardest map solved in its 60 certified moves. In the memory task, training used 9 of 35 combinations of delay and input timing. We scored each timing by net memory use: 1 when the model solves the histories of both cues, 0 when it solves only one, as a model that ignores the cue would, and *−*1 when it solves neither. We trained four learners: the readout alone or the brain and readout in scene 1, and the readout alone in scenes 2 and 3, two further board layouts of the task. In all 34 replicates of the four learners, the model uses the cue at the trained timings and not at the others (Holm-adjusted *P* = 0.0039 to 0.016; Figure 1I), and every run replays bit for bit (Figure S1I). We next asked how a network without a dedicated memory circuit holds the cue through the silent delay.

### Working memory is held in an activity-silent intrinsic trace

To find where the cue is held, we recorded every state variable of every cell through single trials (Figure 2A; Video S2). Spikes that tell the two cues apart stop 22 to 42 ms after the cue ends in the four learners (Figure 2B; Figure S2A), yet the cue persists. In the trial of Figure 2A, 2,769 cells hold a difference in spike-frequency adaptation between the cues through the delay (Figure 2C; Figure S2C). Of all state variables, only adaptation keeps the cue, and its difference decays with the adaptation time constant of 200 to 202 ms (Figure 2D; Figure S2D). Holding the trace costs no spikes: in none of the four learners does a cell spike differently during the last 598 ms of the 640-ms delay (Figure 2I; Figure S2H).

**Figure 2.**
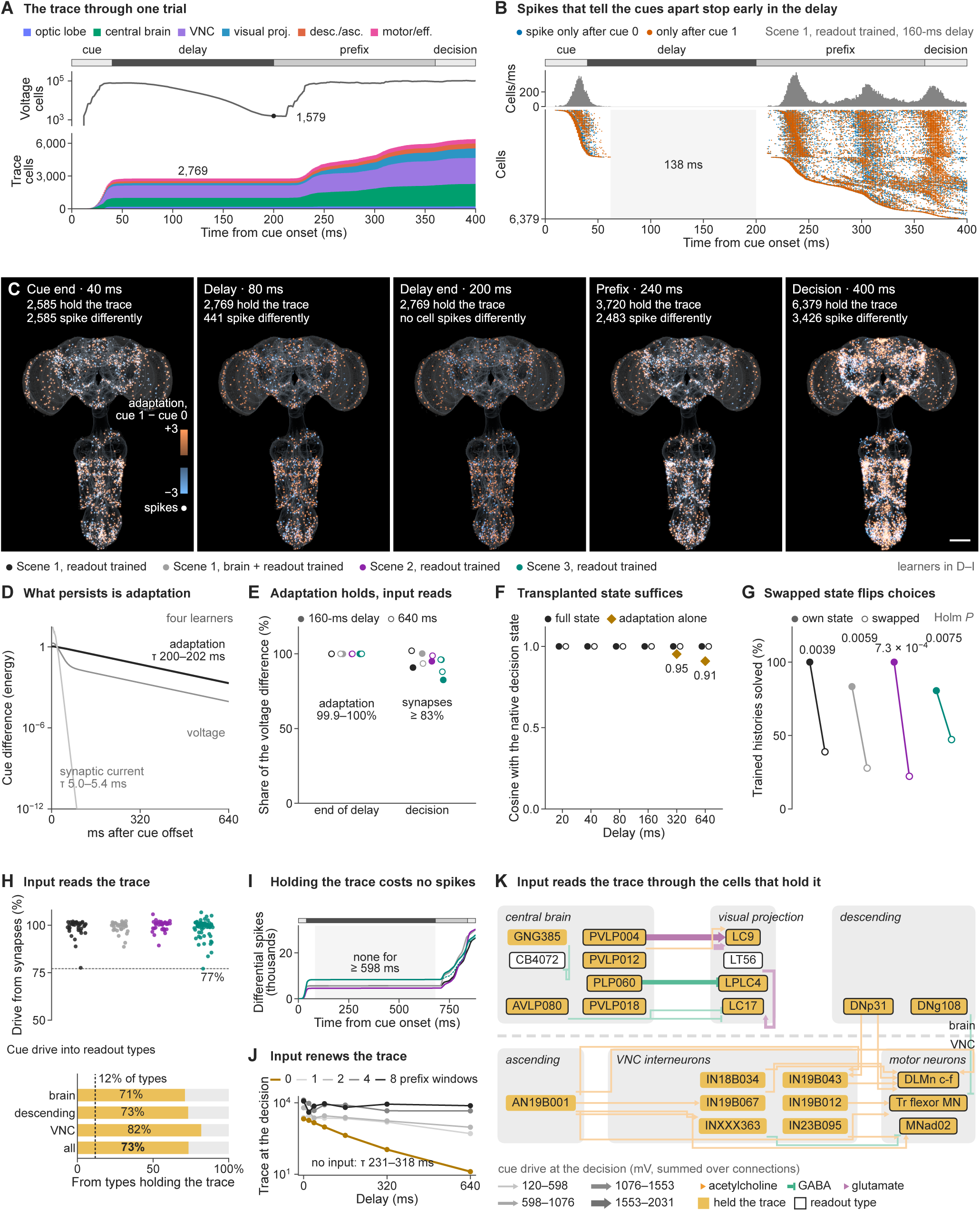
Working memory is held in an activity-silent intrinsic trace. (A) One trial (scene 1, readout trained, 160-ms delay), ms by ms: cells with a voltage difference between the cues (log scale) and cells holding the adaptation trace, by group. (B) Cells that spike only after cue 0 or only after cue 1, with the cells per ms above; shaded, none. (C) The trace (adaptation, cue 1 minus cue 0) at five moments of the trial; white, cells spiking differently. Scale bar, 100 μm. (D) Decay of the cue difference by state variable, relative to early in the delay; lines, exponential fits (time constants over four learners). (E) The share of the voltage difference from adaptation at the end of the delay and from synaptic input at the first decision, four learners, after the 160-ms (filled) and 640-ms (open) delay. (F) Transplants of the full state or of adaptation alone at the end of the delay: cosine with the native decision state. (G) Trained histories solved with the own or the other cue’s state, all trained timings, four learners; exact tests, Holm-adjusted. (H) Top, the synaptic share of the decision-time drive at all 35 timings, four learners; dashed, the minimum. Bottom, the cue drive into readout types from types that held the trace, by region; dashed, their share of types. (I) Cumulative differential spikes through the 640-ms delay, four learners. (J) The trace at the decision by delay and prefix input; the line without input is the fitted decay. (K) The cue drive at the first decision between cell types of the brain (above) and ventral nerve cord (below): width, drive; color and arrowhead, the sender’s transmitter; gold, types that held the trace; outlined, readout types. See also Figure S2 and Video S2.

The next input reads the trace out. At the end of the delay the cue difference in membrane voltage comes from adaptation, but at the decision it comes from synaptic input (Figure 2E), and at least 77% of the decision-time drive is synaptic at all 35 timings (Figure 2H; Figure S2E). Input also renews the trace, which otherwise decays as fitted (Figure 2J).

The trace is sufficient and specific. Moving the full state at the end of the delay into another trial recreates the native decision (cosine 1.000; Figure 2F; Figure S2F). Swapping the states of the two cues flips the decisions at all trained timings in all four learners (Holm-adjusted *P* = 7.3 × 10^−4^ to 0.0075; Figure 2G). The input reads the trace through the cells that hold it: the 12% of cell types that held the trace supply 73% of the cue drive into readout types, in the brain and the ventral nerve cord alike (Figures 2H and 2K). The cue at the decision is spread over many cells in every learner (Figure S2B). We next asked where in the CNS the trace sits, and whether any circuit is special.

### The trace is distributed, and no circuit is special

Memories are often assigned to ensembles of cells. ^27^ We took the 1% of cells that carry the most cue difference at the decision as the cue ensemble of each of two scenes (1,672 cells in scene 1; Figure 3A; Figure S3A). The ensemble is densely wired, with 1.7 and 2.5 times the internal connections of 20 matched sets in the two scenes (Figure 3B), and it is made of the same cell groups as the rest of the CNS (Figure S3B).

**Figure 3.**
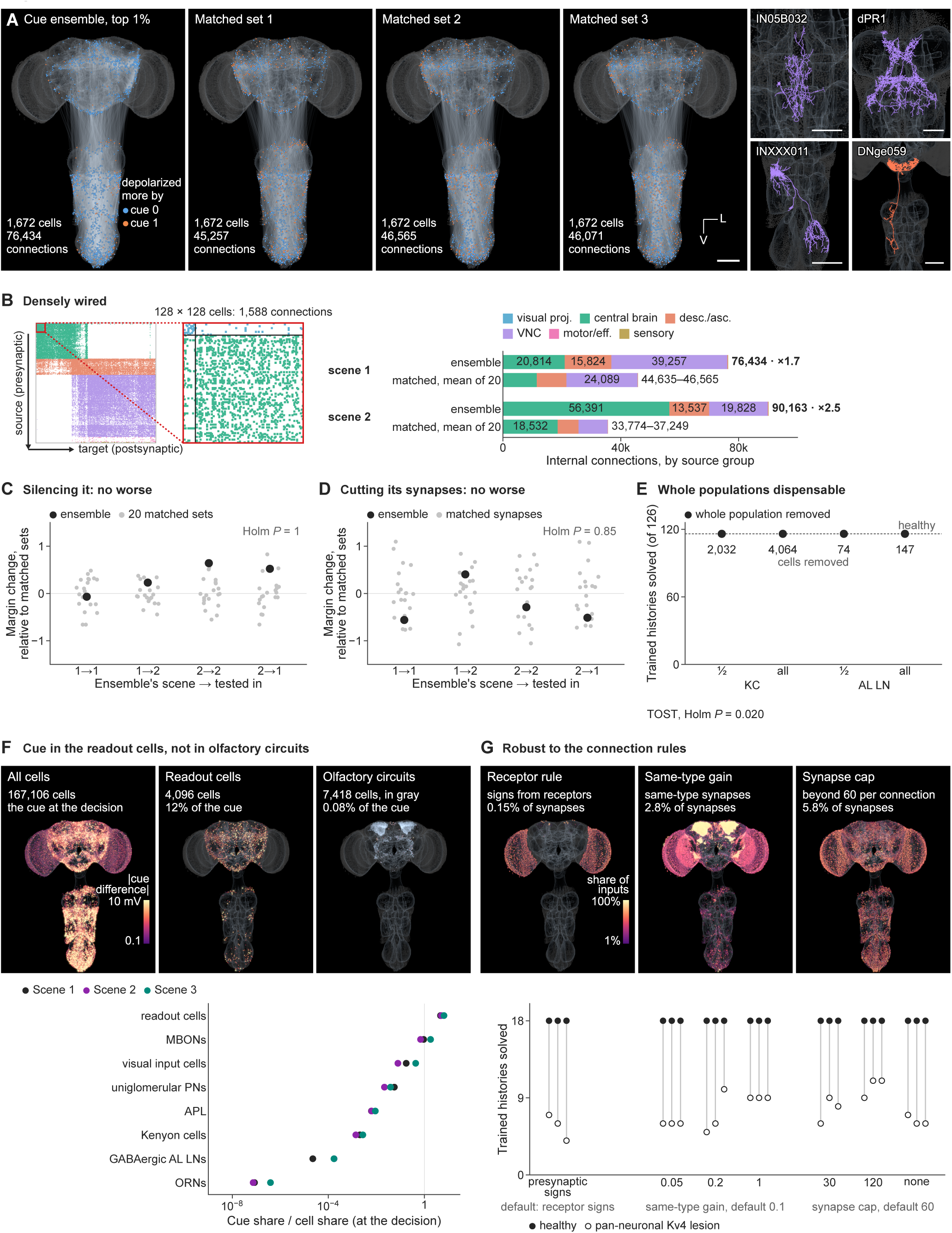
The trace is distributed, and no circuit is special. (A) The scene-1 cue ensemble (the 1% of cells with the largest cue difference at the decision) beside three of its 20 matched sets, which share its cell types, distance from the input, degree and readout role; right, four cue-tuned cell types. Scale bars, 100 μm and 50 μm (cell types). (B) The ensemble’s connections as a matrix (sources down, targets across) with its densest block enlarged, and its internal connections by source group against the mean of its 20 matched sets, two scenes. (C) Silencing the ensemble at the decision: the change in decision margin relative to the matched sets’ mean (gray, each matched set), ensembles of two scenes tested in both; one-sided tests, Holm-adjusted. (D) Removing the ensemble’s internal synapses, against matched synapses, as in (C). (E) Whole populations removed: trained histories solved of 126; dashed, healthy; equivalence tests (TOST), Holm-adjusted. (F) Maps of the cue difference at the decision, and each circuit’s share of the cue against its share of cells, three scenes. (G) Where three connection rules act, and trained histories solved under each variant rule (three runs each), healthy (filled) and under the pan-neuronal Kv4 lesion (open). See also Figure S3.

Yet the ensemble is not necessary. Silencing it at the decision changes the decision no more than silencing matched sets (Holm-adjusted *P* = 1.00; Figure 3C), and cutting its internal synapses does no more than cutting matched synapses (*P* = 0.85; Figure 3D; Figure S3C). Whole populations are dispensable: with half or all of the Kenyon cells or the antennal-lobe inhibitory neurons removed, the network solves the trained histories that the healthy network solves, 116 of 126 (equivalence test, Holm-adjusted *P* = 0.020; Figure 3E; Figure S3D).

The cue concentrates where it is read, not in olfactory circuits: the readout cells carry 12% of the cue difference, 4.7 to 6.6 times their share of cells (Figure 3F). None of this depends on how the connectome becomes a network. All 24 runs learn under 8 sets of connection rules, and the signatures of healthy circuits hold under each (Figure 3G; Figure S3E). If no circuit is special, the cells that hold the trace might still share a property, and we asked what marks them.

### Trace-carrying cells have a molecular signature: *SK*

To ask what marks the cells that hold the trace, we joined published single-cell atlases of the fly brain and ventral nerve cord ^48–53^ to the connectome, by cell type and, in the ventral nerve cord, through 21 hemilineages (Figures 4A–4C; Figures S4A and S4B). The join reproduces the atlas clusters (Spearman ρ = 0.82) and the fast transmitter predicted from electron microscopy ^54^ in 21 of 21 hemilineages (Figure 4H; Figures S4C–S4E). It maps 32% of the trace carriers (the cells that hold the trace at the first decision), up from 7% by cell type alone, and 54% in the ventral nerve cord (Figure 4F).

**Figure 4.**
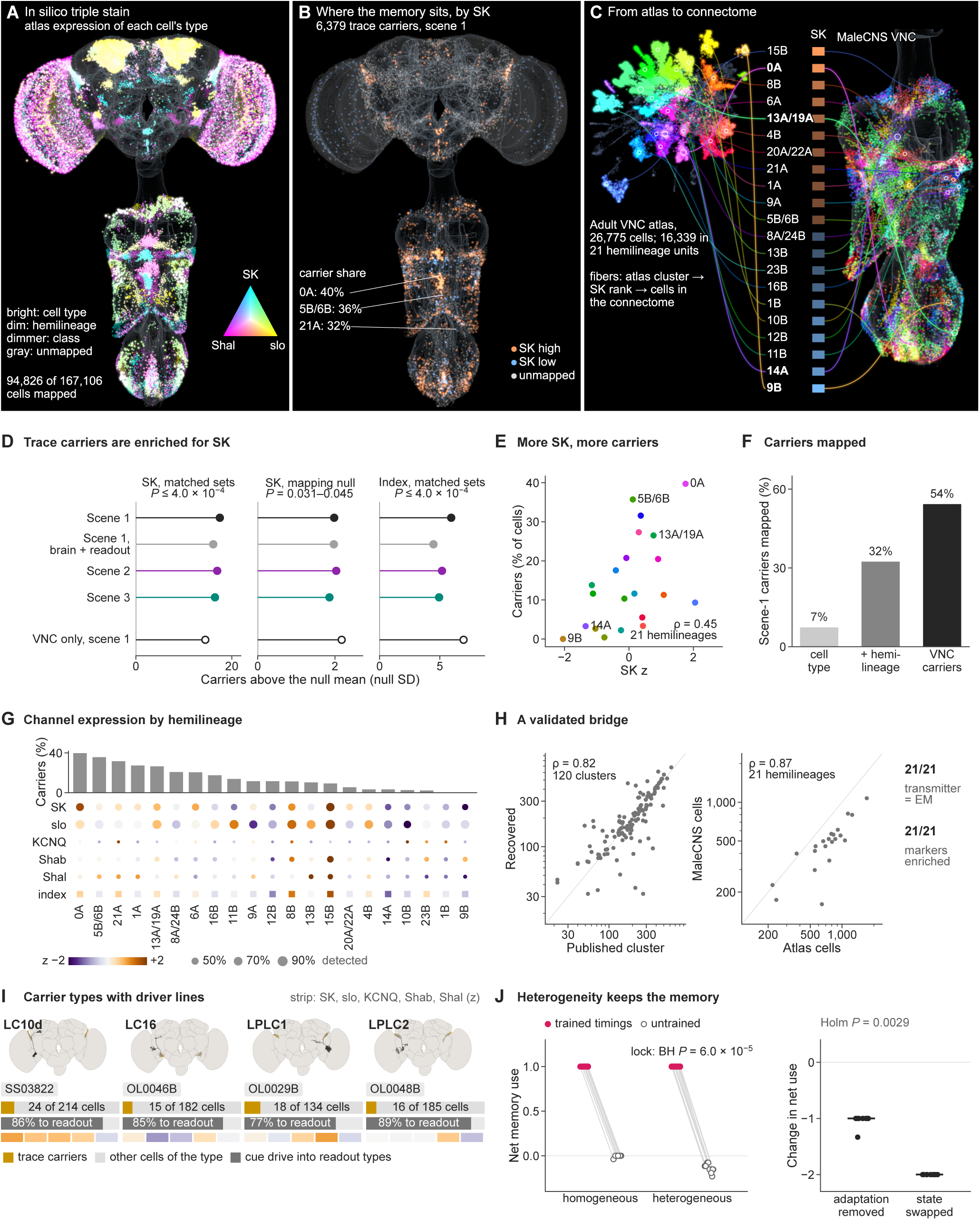
Trace-carrying cells have a molecular signature: *SK*. (A) Three channel transcripts (*SK*, *Shal*, *slo*) painted on the connectome from each cell’s atlas unit: bright, cell type; dim, hemilineage; dimmer, class; gray, unmapped. (B) The trace carriers of scene 1 (cells holding the trace at the first decision), by *SK*. (C) The ventral nerve cord atlas joined to the connectome through 21 hemilineages, ordered by *SK*. (D) Carrier enrichment in *SK* and in the adaptation index, against matched sets and a mapping-shuffle null, four learners and the ventral nerve cord alone (open), in null SD; adjusted *P* from the declared families. (E) Carrier share against *SK* across 21 hemilineages. (F) Scene-1 carriers mapped by cell type, with hemilineages, and in the ventral nerve cord. (G) Carrier share and the channel panel across hemilineages: color, z score; size, share of cells detected. (H) Validation of the bridge: recovered against published cluster sizes, connectome against atlas cells per hemilineage, and the transmitters and markers that agree. (I) Carrier types with split-GAL4 lines: each type on the brain, its carriers among its cells, the share of its cue drive into readout types and its channel strip. (J) Brains with homogeneous or atlas-informed heterogeneous adaptation, 10 runs each: net memory use joined by run, and the change when adaptation is removed or states are swapped; bars, medians; exact tests, Holm-and BH-adjusted. See also Figure S4.

As we declared before testing, the carriers are enriched for *SK*, which encodes a small-conductance calcium-activated potassium channel. Carriers exceed matched sets in *SK* and in an adaptation index (adjusted *P* ≤ 4.0 × 10^−4^), and *SK* exceeds a null that shuffles the mapping (adjusted *P* = 0.031 to 0.045; Figure 4D; Figure S4F). The ventral nerve cord, whose data never shaped the prediction, confirms it: across 21 hemilineages, the share of carriers rises with *SK* (Spearman ρ = 0.45; Figure 4E), against the background of the full channel panel (Figure 4G).

The carriers are accessible in the fly. Four carrier cell types with split-GAL4 driver lines send 77% to 89% of their cue drive into readout types (Figure 4I), and the cue reaches the motor neurons through *SK*-rich hemilineages (Figure S4G). Adaptation that varies from cell to cell, as the atlases suggest, leaves the mechanism intact: brains with atlas-informed heterogeneity keep the memory, its lock to the trained timings and the silent trace in all 10 runs (Holm-adjusted *P* = 0.0029; Figure 4J). If the trace is the memory, something else must decide when it can be read.

### Access to the memory is locked to the timing of learning

The readout uses the trace only at the timings it was trained on. Net memory use is 1 at all 9 trained timings in every replicate and averages −0.004 at the other timings (Holm-adjusted *P* = 0.0039; Figure 5B; Figure S5B), and the lock holds in 66 of 66 runs across nine scenes (Figure 5C).

**Figure 5.**
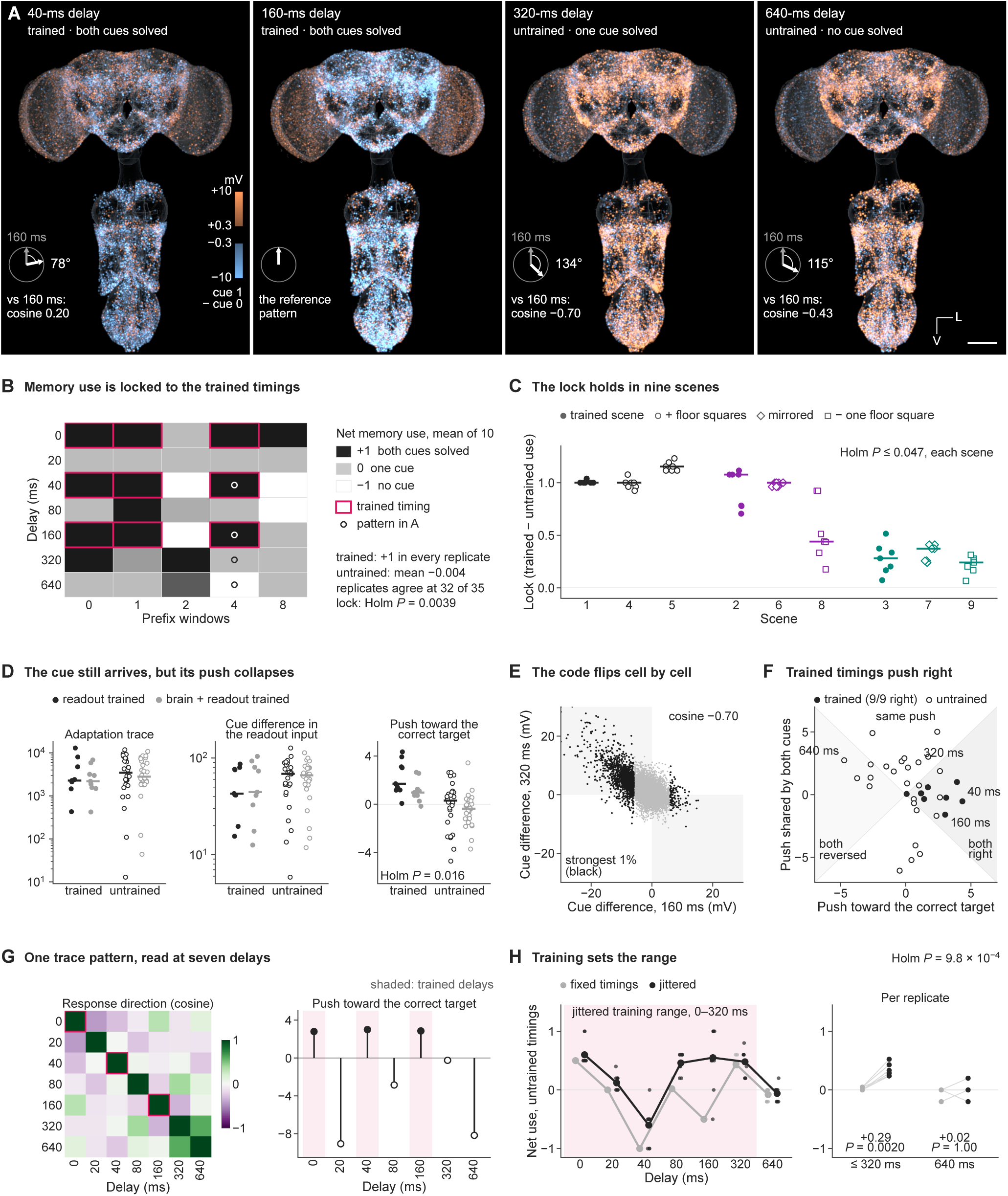
Access to the memory is locked to the timing of learning. (A) The cue difference in membrane voltage at the first decision after four delays, with each pattern’s angle and cosine to the trained 160-ms pattern. Scale bar, 100 μm. (B) Net memory use on the grid of delay and prefix input, mean of 10 replicates; outlined, trained timings. (C) The lock (trained minus untrained use) per run in nine scenes, the trained scenes and their variants; bars, medians; exact sign-flip tests, Holm-adjusted. (D) Trained against untrained timings, two learners: the adaptation trace at the decision, the cue difference in the readout input and the push toward the correct target; bars, medians; exact tests, Holm-adjusted. (E) Each cell’s cue difference after 160 against 320 ms; black, the strongest 1% of cells. (F) Decision geometry at every timing: the push toward the correct target against the push shared by both cues. (G) One trace pattern injected at seven delays: the cosine between response directions, and each response’s push; shaded, trained delays. (H) Training at fixed or jittered timings: untrained use by delay (lines, means of 10 replicates; dots, replicates), and per replicate inside and outside the jittered range; two-sided sign-flip tests, Holm-adjusted. See also Figure S5 and Video S3.

The lock is not a loss of the trace. At untrained timings the trace reaching the decision is no smaller (median size 3,461 against 2,279 at trained timings), but its push toward the correct target collapses (Holm-adjusted *P* = 0.016; Figure 5D; Figure S5C). After the untrained 320-ms delay, the decision-time pattern is nearly the mirror image of the trained 160-ms pattern (cosine −0.70; Figure 5A; Figure S5A), and 92% of the strongest cells change sign (Figure 5E; Figure S5D). In the space of decisions, trained timings push toward the correct target, and untrained timings do not (Figure 5F).

The cause is a drift in how the trace is read. Injected at seven delays, one fixed trace pattern pushes toward the target only at the trained delays (+2.8 to +3.0; Figure 5G; Figure S5E). The operator that turns the trace into a push therefore changes with delay while the trace persists, and we call this operator drift. Every cell group turns with it (Figure S5D). Training across jittered timings widens use inside the jittered range (+0.29) but not beyond it (+0.02; Figure 5H; Figure S5F), and Video S3 shows one trace read at a trained and an untrained delay. We next asked whether access is also locked to the network’s excitability.

### The wiring is robust; access follows the trained states

The connectome keeps the readout within reach. With 90% of the other cells removed at random, 99.85% of the readout cells remain reachable from the visual input, while the cells on routes fall from 116,326 to 11,219 (Figures 6A and 6B; Figures S6A and S6B). Routes thin through every cell class, but none breaks: three-step routes fall to 0.40 of intact, and 3,060 of 10,185 cell types still carry them (Figures 6G and 6H; Figure S6G).

**Figure 6.**
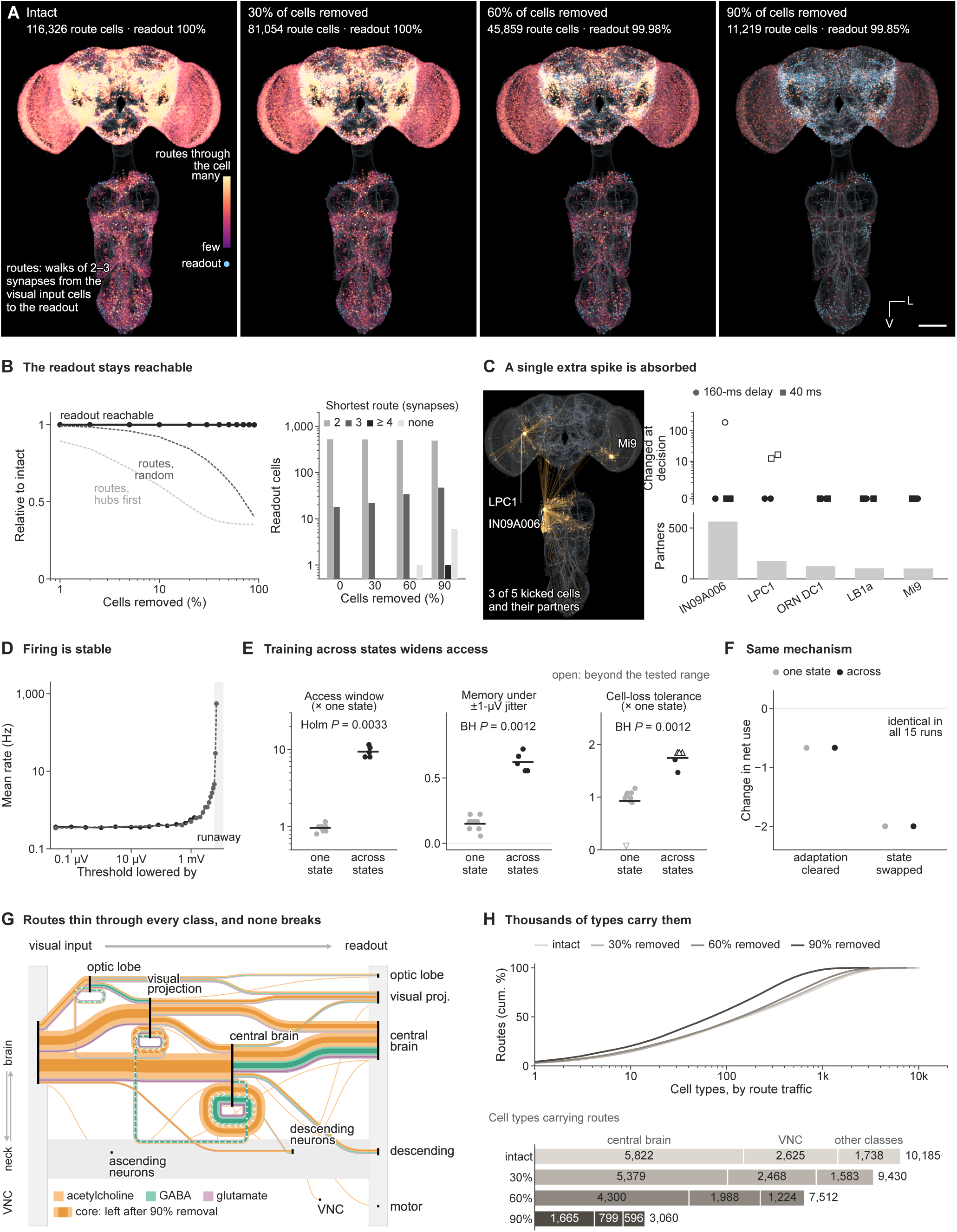
The wiring is robust; access follows the trained states. (A) Cells on routes (walks of two or three synapses from the visual input to the readout), intact and after random removal of 30%, 60% and 90% of the other cells; color, routes through each cell. Scale bar, 100 μm. (B) Readout cells reachable and routes kept under random or hubs-first removal, and the readout cells by shortest route. (C) Single extra spikes in five cells at two delays: three kicked cells with their partners, and the cells changed at the decision. (D) Mean firing rate against a uniform lowering of the threshold; shaded, runaway. (E) Training across threshold states (5 runs) against training at one state (10): access window and cell-loss tolerance relative to the median at one state, and memory under held-out threshold jitter. Bars, geometric means (window, tolerance) and means (memory); open, beyond the tested range; exact tests, Holm-and BH-adjusted. (F) The same mechanism in both: the change in net use when adaptation is cleared or states are swapped. (G) The route network by cell class: strands by transmitter; darker cores, the routes left after 90% removal. (H) The cumulative share of routes across cell types ranked by traffic, and the cell types carrying routes at each level, by class. See also Figure S6.

The dynamics are stable. A single extra spike leaves nothing at the decision in 17 of 20 kicks (Figure 6C; Figure S6C), and the mean firing rate stays between 0.35 and 0.56 Hz for threshold changes of up to 1 mV (Figure 6D).

Access follows the states of training. Decoder theory predicts that a readout trained across excitability states keeps working where one trained in a single state fails, ^36^ and the model bears this out. Training across threshold states widens the access window 9.8-fold (95% interval 8.5 to 11.3; Holm-adjusted *P* = 0.0033). It also keeps memory under threshold jitter (0.62 against 0.15) and raises tolerance to cell loss 2.2-fold (1.6 to 3.9; Figure 6E; Figures S6D and S6E). The mechanism is the same in both: a silent adaptation trace read by the input (Figure 6F). We then asked what happens when disease shifts the state after learning.

### Disease steals access, not memories

We screened 24 fly pathologies for credible operators, requiring published evidence, a mechanism the model can express, healthy circuits that match the fly and the fly phenotype reproduced in two brains (Figure 7A; Figure S7B; Table S1). Of these, 4 passed, and only one reaches visual memory: the pan-neuronal loss of the A-type potassium channel Kv4 that Aβ42 causes in flies ^46^ (Figure 7F).

**Figure 7.**
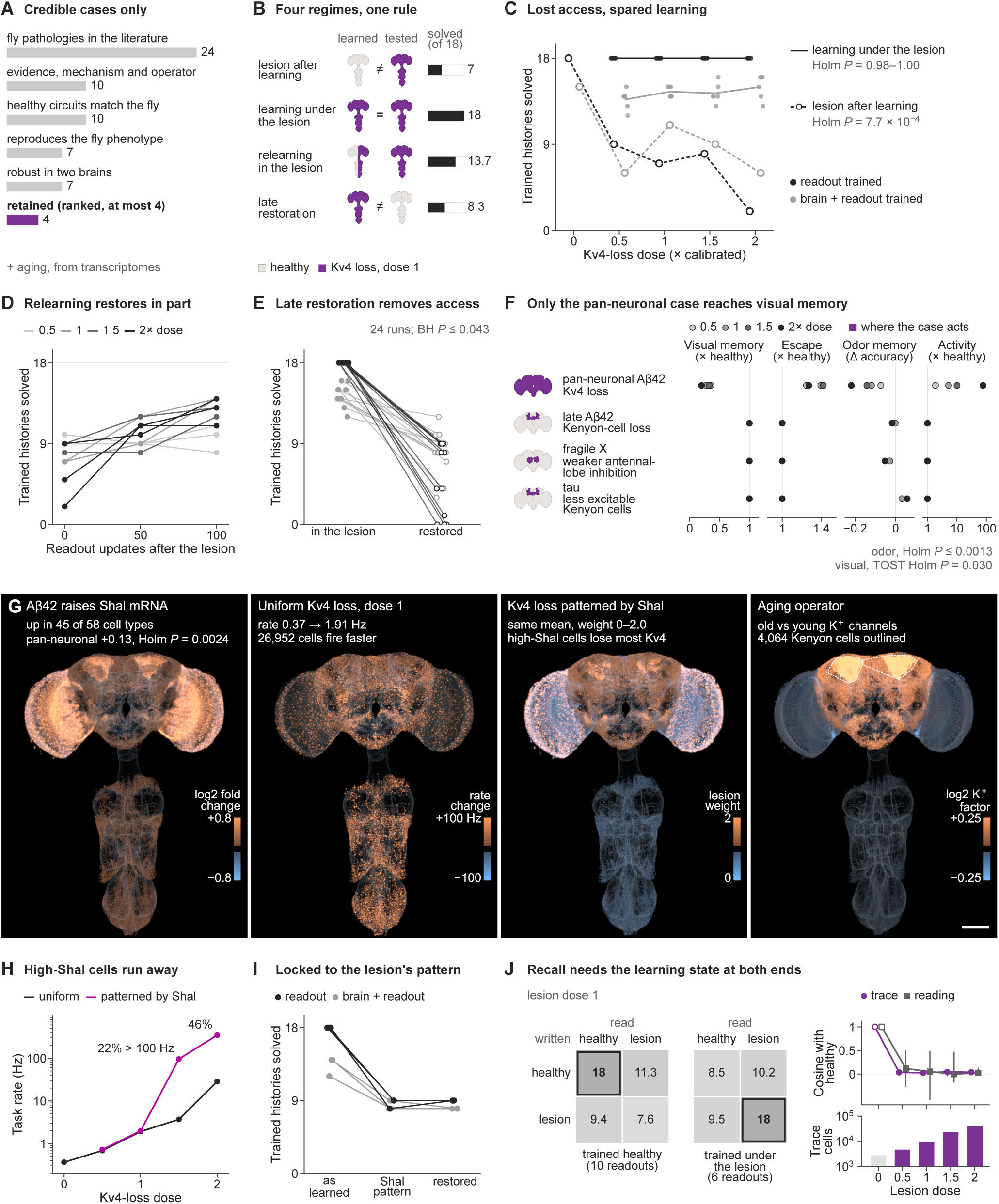
Disease steals access, not memories. (A) The credibility screen of fly pathologies (Table S1). (B) Four regimes: the brain state at learning and at test, and the trained histories solved of 18 (dose 1, readout trained). (C) Trained histories solved after a lesion and when learning under it, by dose, two learners; lines, means; Page trend tests, Holm-adjusted. (D) Relearning after the lesion: trained histories solved after readout updates, by dose (12 runs). (E) Late restoration of excitability after learning under the lesion (24 runs); exact tests, BH-adjusted. (F) Phenotype profiles of the four credible cases by dose, and where each acts; comparison and equivalence (TOST) tests, Holm-adjusted. (G) Disease on the brain: *Shal* transcripts under Aβ42, firing under the uniform Kv4 loss, the Kv4 loss patterned by *Shal*, and the aging operator. Scale bar, 100 μm. (H) Task firing rate under the uniform and the *Shal*-patterned Kv4 loss, by dose. (I) Trained histories solved by learners trained under the uniform lesion: as learned, under the *Shal* pattern and with excitability restored. (J) Recall needs the learning state at both ends. Left, trained histories solved when the trace is written in one state and read in another (the full state moved at the end of the delay), dose 1; means of readouts trained healthy (10) or under the lesion (6). Right, across doses, the cosine with the healthy brain of the stored trace and of the healthy trace’s reading (median and range over the trained timings), and the cells holding the trace (median). See also Figure S7 and Video S4.

A lesion after learning removes access, whereas learning under it is spared (Figure 7B). At dose 1, the strength that reproduces the fly phenotype, the Kv4-loss lesion lowers the trained histories that the learned readout solves from 18 to 7, and memory falls with dose (Holm-adjusted *P* = 7.7 × 10^−4^; Figure 7C; Figure S7C). Readouts trained in the lesioned brain solve 18 of 18 in every one of 48 runs (Figure 7C; Figure S7D). Retraining the readout after the lesion recovers 0.41 of the loss (Figure 7D; Figure S7E), but restoring excitability late, after learning under the lesion, removes access again and leaves 8.3 of 18 (BH-adjusted *P* ≤ 0.043; Figure 7E; Figure S7F). Memories learned under one lesion pattern are likewise lost under another (Figure 7I).

An atlas of Aβ42 flies constrains how the lesion acts: neuronal *Shal* transcripts rise in it (+0.13 log_2_), ^52^ so the Kv4 loss must arise after transcription (Figure 7G). Patterned by *Shal*, the lesion drives 22% to 46% of cells above 100 Hz at 1.5 to 2 times the dose (Figure 7H; Figures S7H–S7J).

To ask whether the lesion damages the stored trace or its reading, we moved the full state of each trial at the end of the delay between the healthy and the lesioned brain (Figure 7J; Video S4). Recall needs the learning state at both ends. Readouts solve all 18 trained histories when the trace is written and read in the state of learning, and at most 12 when either end differs. At dose 1, a healthy trace read under the lesion gives 11.3 of 18 and a lesion-written trace read in health 9.4; for readouts trained under the lesion, these are 10.2 and 9.5.

The lesion changes both ends. It writes the trace into 4,683 to 39,285 cells instead of 2,769, nearly orthogonal to the healthy trace (cosine 0.03 to 0.05). The hold stays silent only at the lowest dose: 0 cells spike differently late in the delay at dose 0.5, against 11,240 at dose 1. Read under the lesion, the same healthy trace evokes a decision-time pattern nearly orthogonal to its healthy reading (median cosine 0.00 to 0.12). The lesioned brain also spends more for less: across doses, a trial costs 1.9 to 77 times the spikes of a healthy trial for 0.01 to 0.04 bits about the cue, against 0.73 bits in health (Figure S7I). An aging operator built from transcripts of the aging fly ^51^ dims Kenyon-cell odor responses with its strength (Figure S7K). In the model, memory failure is a change of state at writing and at reading, not a loss of storage.

## DISCUSSION

A fixed connectome, trained end to end, holds working memory in neuronal adaptation and recalls it only in the brain state that wrote it. The trace needs neither persistent firing nor a dedicated circuit, and the learned readout can use it only where the operations that write and read it match the state of learning. The result joins two questions that are usually asked apart: what holds a silent memory, and why recall depends on the state of the brain.

One dependence accounts for four observations in the model. The operator that turns the silent trace into a push depends on the state of the network, in time and in excitability. Hence the lock to the trained timings, its widening when training spans timings or excitability states, the loss of access after a lesion and its partial return on relearning. Excitability-driven decoder drift ^36^ and state-dependent retrieval ^34,35^ then become two sides of one rule: recall spans the states of learning, at writing and at reading. Context-dependent memory in humans, with recall best in the setting of learning, ^55^ may express the same rule at the scale of whole environments. Because the lesion rewrites the trace as well as its reading, restoring either end alone does not restore recall.

The wiring itself is robust. Routes from the input to the readout persist after heavy cell loss, and a single extra spike is absorbed before the decision. What fails under the lesion is therefore not reachability but the match between the state that wrote a memory and the state that reads it. In animals, neuromodulators shift the intrinsic properties on which such a match rests: cholinergic activation, for example, switches the firing mode of hippocampal CA3 neurons. ^56^ Training across states widens the match, and cell-to-cell differences in adaptation leave the mechanism intact, in line with the robustness that heterogeneity lends to learning in spiking networks. ^57^ Varied conditions during learning may therefore protect a memory against later shifts of state.

Learning on real wiring recruits adaptation, an intrinsic and cell-autonomous variable, rather than persistent firing or synaptic facilitation. ^19–21^ Holding the cue costs no extra spikes, an economy that matters because neural signaling is energetically expensive. ^58^ The time dependence of the readout also has a concrete place. In primate recordings, stable population codes sit beside impulse responses that change with time; ^30,31^ in the model, the trace persists through the delay while the operator that reads it drifts, so the time dependence lies in the reading, not in the code. Models of representational drift ask how stable readouts can track a code that keeps changing; ^59^ the model poses the converse problem, a persistent code read by a changing operator, which the learned readout solves only over the timings it was trained on.

The carriers of the trace are enriched for *SK* in the fly’s own transcriptomes. SK currents shape after-hyperpolarization and sparse coding in fly circuits, ^60^ and enhancing them impairs hippocampal memory in mice. ^61^ The signature sides with a neurocentric view of memory ^62^ without implying that flies hold working memories in SK currents: the enrichment marks where to look. The delays here last under a second, shorter than the visual memories of seconds that depend on signaling in ring neurons of the fly’s central complex, ^24,25^ so the two mechanisms may serve different time scales.

No circuit in the model is necessary for the memory. Its strongest ensemble is densely wired yet no more necessary than matched sets, and removing all Kenyon cells, the intrinsic cells of the mushroom body, leaves visual memory intact. Flies without mushroom bodies likewise keep visual learning, ^63^ although mushroom-body circuits can also store visual memories. ^64^ Engrams of a single memory are spread across many brain regions in mice; ^27^ in the model, the spread follows from where the trace is written and read, and the cue concentrates in the readout cells, several times out of proportion to their number. Excitability biases which neurons join an engram, ^28,29^ and adaptation currents carried by SK channels may set that bias for a silent working memory.

The model also bears on when to treat. A disease-like shift after learning rewrites the trace and changes its reading, so normalizing excitability late is a second shift, one that removes the access learned under the disease. Restoring excitability should therefore help new learning more than the recall of memories formed during the disease. This reading is consistent with the mixed effects of lowering hyperexcitability in patients, ^43–45^ with the recovery of memories by direct engram activation in early disease models ^37^ and with the rescue of retrieval by activating parvalbumin interneurons in such mice. ^38^ Because hyperexcitability emerges long before diagnosis, ^41,42^ a progressing disease keeps moving memories away from the state in which they were formed, and treatments that hold the state steady early may protect access better than treatments that reverse it late.

Connectome models have so far predicted pathways and activity without learning, or learned within one sensory pathway. ^6,7^ A digital twin trained on tasks allows experiments that no animal allows, such as moving a stored trace from one brain state into another, and each of its results is a prediction for the fly. Because every experiment here was declared before it ran and replays bit for bit, others can re-examine the twin with the same certainty. Coupled to a physics model of the fly’s body, ^9^ a trained twin could also link memory to movement in a closed loop.

Six of these predictions can be tested directly. Knocking down *SK* in carrier hemilineages (0A, 13A/19A, 6A and 21A), which driver lines can target, ^65^ should shorten visual delay memory more than knocking it down in *SK*-poor ones (9B, 14A and 10B). An impulse during the delay should reveal the cue in the cell types the model marks. ^15^ Training across sustained excitability shifts too small to change behavior should protect later recall, whereas a shift confined to encoding, or to recall, should impair short visual memory. With temporal control of transgene expression, ^66^ memories formed before Aβ42 is induced should be lost while new learning works, ^46^ and acute restoration of Kv4 after training under Aβ42 should impair that memory. Memory failure, in this model, need not mean lost storage: it can mean a key that no longer fits its lock.

### Limitations of the study

Our results come from a model, so each statement about flies or patients is a prediction that awaits a test in the animal. We built the model on the connectome of one male fly, and some cell types and connections differ between the sexes. ^2,5^ Female connectomes of the brain and of the nerve cord came from separate animals. ^1,67^ A connectome that unites the brain and nerve cord of one female fly ^3^ will allow the same training and tests in the other sex, once its proofreading and cell typing match those of the male CNS. Each connectome is also a snapshot of one animal: cell types recur across individuals, with occasional variation in neuron counts and connections. ^4^ Twins trained on several connectomes would separate what is general in the mechanism from what is individual.

During the task, synapses change only by short-term depression at olfactory terminals, the model has no neuromodulation, and its delays last under a second, shorter than the visual memories of several seconds in behaving flies. ^24,25^ Adding short-term synaptic dynamics across the CNS and the dopamine-gated plasticity that writes associative memories in the mushroom body ^68^ would show whether the intrinsic trace competes with synaptic memories over longer delays or complements them. The transplant between brain states moves the full state, including its cue-independent part, so it separates where a trace was written from where it was read, not the trace from the state that carries it. Transplants of single state variables can make that separation.

The molecular links rest on transcripts. Channel transcripts can track channel currents in identified neurons, ^69^ but disease can uncouple the two: in Aβ42 flies, *Shal* transcripts rise ^52^ while the Kv4 channel is degraded. ^46^ The atlases map 32% of the trace carriers, and most of the others lie in the brain, where atlases that resolve lineages ^70^ or span the whole body ^71^ can extend the join. The disease operators are calibrated to published fly phenotypes rather than measured in the model’s own cells. Recordings from identified neurons of Aβ42 flies would calibrate them cell by cell, and the twin could then follow how such shifts reshape memory as a disease progresses.

## RESOURCE AVAILABILITY

### Lead contact

Requests for further information and resources should be directed to and will be fulfilled by the lead contact, Chunjie Tian.

### Materials availability

This study did not generate new unique reagents.

### Data and code availability

- This paper analyzes existing, publicly available data, accessible at the DOIs and accession numbers listed in the key resources table. The simulation records, trained models and result tables have been deposited at Zenodo as a core record (https://doi.org/10.5281/zenodo.23629827) and an extended record (https://doi.org/10.5281/zenodo.23629835). The files are under restricted access during peer review. Editors and reviewers may obtain access from the lead contact. A manifest lists every deposited file with its SHA-256 hash.
- All original code is publicly available on GitHub at https://github.com/Hanziwww/fly-connectome-silent-memory, release v1.0.0.
- Any additional information required to reanalyze the data reported in this paper is available from the lead contact upon request.

## ACKNOWLEDGMENTS

This work was supported by the National Natural Science Foundation of China (grant nos. 82160215, 81660174, and 82271169); the Guangdong Provincial Key Laboratory of Autophagy and Major Chronic Non-communicable Diseases (grant no. 2022B1212030003); the Research Startup Funds for High-level Talents in the Affiliated Hospital of Guangdong Medical University (grant no. GCC20220013); the College Students’ Innovative Entrepreneurial Training Plan Program (grant no. S202510571025); and the 2025 Artificial Intelligence-Enabled Medical Device Innovation Task (Open Competition Mechanism, project no. 2025-AISJK-02007).

## AUTHOR CONTRIBUTIONS

Han Wang, Joe Yeong Poh Sheng, and Chunjie Tian conceived and designed all studies. Han Wang wrote all code, performed all analyses, generated all figures and videos, and prepared all supplemental materials. Joe Yeong Poh Sheng contributed to writing the manuscript. The other authors reviewed and revised the manuscript. All authors reviewed and approved the final manuscript and take responsibility for its content.

## DECLARATION OF INTERESTS

The authors declare no competing interests.

## DECLARATION OF GENERATIVE AI AND AI-ASSISTED TECHNOLOGIES IN THE MANUSCRIPT PREPARATION PROCESS

During the preparation of this work, the authors used Claude (Anthropic) solely for language review. The authors reviewed and edited the manuscript and take full responsibility for its content.

## SUPPLEMENTAL INFORMATION

**Document S1.** Figures S1–S7 and Table S1.

**Table S2. Every declared test the paper reports, related to Figures 1–7**. Each test with its family, where the paper reports it, its statistic and n, the estimate, *P*, *P* adjusted within the complete declared family, the family’s size and correction, and whether it is primary or post hoc (Excel).

**Table S3. The run registry, related to Figures 1–7**. Every training and evaluation run with its configuration, its record and its outcome (Excel).

**Table S4. The single-cell panels, related to Figure 4**. The channel and marker genes with their FlyBase identifiers, and their values in every atlas unit (Excel).

**Data S1. Source data for Figures 1–7 and S1–S7.** The values behind every panel, one table per figure (.csv).

**Video S1. The trained connectome plans, move by move, related to Figure 1**. The final brain and readout solve the hardest planning map in its 60 certified moves. Left, the board and the readout’s choice among the four moves (the softmax of its logits; the chosen move white). Right, the unfolded CNS, with every cell that spikes in the move’s 40-ms run in its group color. Below, live charts of the cells spiking per move, stacked by group, and of the readout’s confidence in its move, with box pushes marked.

**Video S2. Write, hold, read: one memory trial, millisecond by millisecond, related to Figure 2**. Scene 1, readout trained, 160-ms delay with four input windows. Left and center, the cells spiking after cue 0 and after cue 1; right, the trace (adaptation, cue 1 minus cue 0) with the cells spiking differently in white; small boards show what the model sees. Live charts show, per ms, the cells spiking differently and the cells holding the trace, and the trace’s strength against the adaptation’s own decay (dashed), which it follows exactly once no cell spikes differently.

**Video S3. One trace, two readings, related to Figure 5**. The trained 160-ms and the untrained 640-ms delay side by side, phase by phase, the longer delay played faster. Each panel shows the trace with the cells spiking differently. Live charts show the cosine between the two trials’ traces at matching moments, each trace’s strength, and each cue’s push toward its target at the first decision. The end card shows the decision-time patterns after 160, 320 and 640 ms with their cosines to the trained pattern.

**Video S4. Lost access, lost repair, related to Figure 7**. Scene 1, readout trained, lesion dose 1. First the regimes of Figure 7B (healthy, the lesion after learning, learning under the lesion, relearning under it and late restoration), then the full state moved between brain states at the end of the delay, as in Figure 7J (white frame). Each column shows the cue-1 history’s spikes, the brain state at learning and test or at writing and reading, and both cues’ histories on the board, ending in a solved or failed mark. Live charts show the cells spiking per ms and, per cue, the readout’s probability of a correct move at each decision.

## STAR METHODS

### Key resources table

| REAGENT or RESOURCE | SOURCE | IDENTIFIER |
| --- | --- | --- |
| **Deposited data** |  |  |
| MaleCNS connectome, v1.0: neuron annotations, transmitter predictions and connection weights | Berg et al.<sup>2</sup> | https://male-cns.janelia.org/download/ |
| Whole-brain single-cell atlas of the aging fly brain | Davie et al.<sup>48</sup> | GEO: GSE107451 |
| Transcriptomes of visual-system cell types (TAPIN-seq) | Davis et al.<sup>50</sup> | GEO: GSE116969 |
| Single-cell atlas of the developing and adult optic lobe | Özel et al.<sup>49</sup> | GEO: GSE142787 |
| Single-cell atlas of the ventral nerve cord, with its published cluster annotations | Allen et al.<sup>53</sup> | GEO: GSE141807; eLife 54074, figure 1 data 1–2 and figure 4 data 1 |
| Aging Fly Cell Atlas, head | Lu et al.<sup>51</sup> | GEO: GSE218661 |
| Alzheimer's disease Fly Cell Atlas, head | Park et al.<sup>52</sup> | GEO: GSE261656 |
| Simulation records, trained models, result tables and source data | This paper | Core: https://doi.org/10.5281/zenodo.23629827;<br>extended: https://doi.org/10.5281/zenodo.23629835 |
| **Software and algorithms** |  |  |
| FlyBrain 0.1.0, whole-CNS spiking simulator of the MaleCNS connectome | C. Benade; model of Shiu et al.<sup>6</sup> | https://github.com/HEREISCB/flybrain |
| Code of this study: simulation, training, analysis, figures and videos | This paper | https://github.com/Hanziwww/fly-connectome-silent-memory |
| Python 3.12.13 | Python Software Foundation | https://www.python.org |
| PyTorch 2.14.0 (CUDA 13.2) and 2.12.1 (CUDA 13.0) | PyTorch Foundation | https://pytorch.org |
| NumPy 2.5.3 | Harris et al.<sup>72</sup> | https://numpy.org |
| SciPy 1.18.1 | Virtanen et al.<sup>73</sup> | https://scipy.org |
| pandas 3.0.6 | pandas development team | https://pandas.pydata.org |
| Numba 0.67.0 | Numba developers | https://numba.pydata.org |
| Matplotlib 3.11.2 | Matplotlib development team | https://matplotlib.org |
| Scanpy 1.12.4 and anndata 0.13.4 | Wolf et al.<sup>74</sup> | https://scanpy.readthedocs.io |
| leidenalg 0.12.0 and igraph 1.0.0 | Traag et al.<sup>75</sup> | https://leidenalg.readthedocs.io |
| scikit-learn 1.9.1 | scikit-learn developers | https://scikit-learn.org |
| statsmodels 0.15.0 | statsmodels developers | https://www.statsmodels.org |
| Pillow 12.3.0 | Pillow contributors | https://python-pillow.org |
| FFmpeg 6.1 | FFmpeg developers | https://ffmpeg.org |
| Claude (language review only) | Anthropic | https://claude.com/claude-code |

### Experimental model and study participant details

This study used no animals and no human participants. Every experiment was a simulation of a model built from the connectome of the complete central nervous system of one adult male *Drosophila melanogaster*. ^2^ The results therefore describe a male CNS; cell types and connections that differ between the sexes ^2,5^ limit their generalization to females, as discussed in the Limitations of the study.

The single-cell atlases joined to the connectome came from published data, with the sexes each study reports. Davie et al. sampled female and male brains of two strains in similar numbers. ^48^ Davis et al. did not sort flies by sex. ^50^ Özel et al. removed male cells from their adult data. ^49^ The ventral nerve cord atlas of Allen et al. has female and male replicates. ^53^ The Aging and Alzheimer’s disease Fly Cell Atlases include both sexes, ^51,52^ and our analysis of the latter treated sex as a covariate.

### Method details

#### The connectome model

We simulated the CNS with FlyBrain (version 0.1.0), a GPU simulator of the MaleCNS connectome that runs the leaky integrate-and-fire model of the fly brain ^6^ on the whole CNS. The model kept all 167,106 neurons and all 25.6 million stored connections of the connectome (directed cell pairs with their synapse counts); none was pruned. In each step of Δ*t* = 1 ms, the membrane potential *v_i_*, synaptic input *g_i_* and adaptation *a_i_* of neuron *i* (all in mV) were updated as

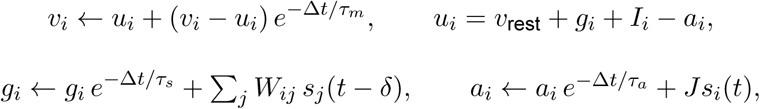

where *I_i_* is an injected current, *s_j_*(*t*) marks a spike of neuron *j* and *δ* is the synaptic delay. A neuron spiked when *v_i_* reached its threshold; it was then reset to *v*_reset_ and held there for the refractory period.

The constants were those of the fly brain model: ^6^ *v*_rest_ = *v*_reset_ = −52 mV, threshold −45 mV, *τ_m_* = 20 ms, *τ_s_* = 5 ms, a refractory period of 2.2 ms, a delay of 1.8 ms (two steps) and 0.275 mV per synapse. Spike-frequency adaptation, which that model lacks, added *J* = 1.5 mV per spike and decayed with *τ_a_* = 200 ms. Short-term depression acted only at the terminals of olfactory receptor neurons and antennal-lobe local neurons (2,952 cells), as at the depressing synapses of olfactory receptor neurons in the fly. ^76^ There, each spike was scaled by the available fraction of a resource and used up 0.2 of it, and the resource recovered with a time constant of 300 ms. Slow metabotropic and monoaminergic currents were off.

The weight of the connection from neuron *j* to neuron *i* was

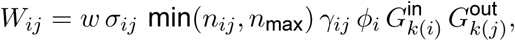

where *w* = 0.275 mV, *n_ij_* is the synapse count, capped at *n*_max_ = 60 synapses, and *σ_ij_* is the sign. The rule gain *γ_ij_*, the fan-in factor *ϕ_i_* and the trained gains *G* of the groups *k* of the two cells are defined below. The sign followed the transmitter predicted for the presynaptic cell; ^2^ where the receptor transcripts of the postsynaptic cell type had been profiled, ^48–50,71,77^ the sign of its dominant receptor class for that transmitter replaced it.

The rule gain was 0.1 between cells of the same cell type and 1 otherwise, with three exceptions. It was 3 from descending neurons onto the ventral nerve cord, 2 from visual projection neurons onto descending neurons, and a further 3 from LC4 and LPLC2 onto the giant fiber DNp01. For neurons with more than 5,000 input synapses, *ϕ_i_* was 5,000 divided by their input synapse count, so that the largest neurons need coherent input to fire.

The board of a task, four channels of 10 by 10 cells (walls, targets, boxes and the player), drove 3,200 visual projection neurons, 8 per channel and cell, each with the board value times 18 mV as a constant input current. The readout observed 4,096 other cells, mostly central-brain intrinsic, descending, visual projection and motor neurons, in 256 groups of 16. Each move was one decision window of 40 ms. The group means of the synaptic input at 14 ms and of the voltage relative to rest at 40 ms gave 512 features, standardized with fixed statistics and clipped at 10 standard deviations. An affine readout mapped them to logits for the four moves, and illegal moves were masked. In the planning task the network state was reset before each decision; in the memory task it persisted through the trial.

#### Training

Training changed only type-level parameters and the readout, never the wiring, the signs or the connection rules. Cells were grouped by cell type, or by superclass where no type was annotated, into 11,770 groups. Each group had an incoming gain and an outgoing gain, both bounded between 0.8 and 1.25, and a threshold offset bounded at *±*1 mV (35,310 parameters in all); the readout had 2,052 weights and biases. Spikes were all or none in the simulation. Gradients passed through a surrogate derivative of the spike, ^78^ 1/(1 + *β|v_i_ − θ_i_|*)^2^ at threshold *θ_i_* with *β* = 25 per mV, and were backpropagated through every step of a trial.

We used Adam with learning rates of 10^−4^ for the brain parameters and 3 × 10^−4^ for the readout, clipped the gradient norms at 1 and 5 and projected the brain parameters back into their bounds after each step. The simulation ran in float64 with gradient sums in float32, the fastest precision that passed a calibration against float64 and float32 on two calibration maps. In the planning task, gradients took two passes: the loss gradient with respect to the readout features, then the brain gradient by recomputing each batch of 16 boards, whose features had to agree with the first pass within 10^−5^. The training seed was 10 throughout.

#### The planning task

The planning task was Sokoban: the model moves the player to push boxes onto targets, and a box pushed into a corner can be lost for good. The ladder held 24 maps in 8 tiers of rising difficulty, three per tier, on boards of 6 to 10 cells across with 1 to 3 boxes. Their optimal solutions, of 9 to 60 moves, were certified by an exhaustive reverse search and checked by an independent forward search. Every map was learned from the same starting model, a brain and readout trained beforehand by the same method on two warm-up maps.

Training followed a reverse curriculum on the certified solution. It began one move before the goal, and each time the model solved the current start on its own, the start moved back by one move; any solution was accepted, detours included. With *m* the margin of the allowed over the disallowed move logits, the loss at the new start was 0.25 log (1 + *e*^(0.10*−*^*^m^*^)/0^.^25^). Decisions already learned whose margin fell below 0.03 added the mean and the maximum of their squared shortfalls, with weight one half each. There were 4 updates per behavioral assessment, and a map was learned when the model solved it from its original start within 512 moves.

Runs in both tasks had wall-clock segments of 20, 40 and 80 min. A run entered the next segment only if it had progressed, by a new curriculum step or by raising its smallest margin by at least 0.02 with every solved step kept; otherwise it stopped by stagnation and kept its best model. Learning cost was the number of updates to learn a map. Besides the original set of maps, the replicate sets (6 in all) differed only in their starting readouts, to which Gaussian noise of 0.1 times the standard deviation of each readout parameter tensor was added. To ask whether brain and readout learned together, we replayed all nine pairings of the brain and readout checkpoints from before, midway through and after training on each map, from its original start.

#### The memory task

In the memory task, the box stands between two targets, one on each side, and a cue shows which of them is in play (Figure 1F). A trial began with a 40-ms cue frame of the board with the cued target. A blank delay without input followed, then input windows of 40 ms that showed the board without targets as the player stepped between two cells, the same for both cues. The model then saw the board without targets, frame by frame, and chose its moves, with the network state carried over from the cue. A trial was solved when the box reached the cued target. It failed when the box was pushed irreversibly toward the other target, when no move was legal or after 512 moves.

The delays were 0, 20, 40, 80, 160, 320 and 640 ms, and the input windows numbered 0, 1, 2, 4 or 8, giving 35 timings. Training used only the delays of 0, 40 and 160 ms with 0, 1 or 4 windows (9 timings); the others were tested only. Each update trained one timing on both cues, cycling through the trained timings. Its loss was the cross-entropy of the certified moves at each decision, with illegal moves masked, averaged over decisions and cues. The gradient ran back through the whole trial, the cue and the silent delay included.

After every 4 updates, free rollouts assessed the model at the trained timings, and the best assessment so far was kept. A run had learned when it solved all trained histories (18 in scene 1) and chose the cued side at the first decision of each. At each timing, net memory use was +1 when the model solved the histories of both cues, 0 when it solved one, as a model that ignores the cue does, and *−*1 when it solved neither. We averaged it over the trained and over the untrained timings of a run; the lock was their difference.

Four learners trained either the readout alone, on the planning-trained brain, or the brain and readout in scene 1, and the readout alone in scenes 2 and 3. Scene 2 is a ring-shaped board, and scene 3 presents one board in two orientations, both learned by one readout. Replicates (10 per scene-1 learner and 7 per scene-2 and scene-3 learner) differed only in the order of the trained timings, a new random permutation for each block of nine updates, and in the readout noise above. Replicate 0 repeated the original protocol, and for three learners it reproduced the original models tensor for tensor.

Six new scenes tested the lock beyond the trained ones. Two were variants of scene 1 with one or two more floor cells; for scenes 2 and 3, the others were the mirror image of each and the first single removal of a floor cell that kept its certified solutions. The readout alone learned each in 7 replicates. Training at jittered timings drew the delay of each update uniformly from 0 to 320 ms in steps of 20 ms, and its input windows from 0, 1, 2, 4, 6 or 8. Each of its replicates shared its starting readout with one fixed-timing replicate of the scene-1 readout learner, and the stopping rule was unchanged.

#### Recording and dissecting the trace

We replayed trained models with every state variable of every cell recorded at every millisecond, for both cues at selected timings. A cell spiked differently at a millisecond when it spiked after one cue and not after the other, and it held the trace when its adaptation differed between the cues. Exponential fits gave the decay of each state variable’s cue difference through the delay. Between spikes the membrane equation is linear. The cue difference in the voltage of a cell that does not spike therefore splits exactly into its decaying initial value, a part driven by synaptic input and a part driven by adaptation; spike resets make up the rest. At the decision, the cue difference of the drive, synaptic input minus adaptation, splits likewise into synaptic and adaptation shares that sum to one.

Transplants replaced, at the end of the delay, the full state of the cue-0 history (voltage, synaptic input and adaptation), or its adaptation alone, with that of the cue-1 history, and compared the decision-time difference that followed with the native one by its cosine. Opposite-cue swaps replaced the full continuous state of every cell at the start of the first decision with the other cue’s state at that moment, at all 9 trained timings; swaps from the same cue changed no decision. For the injections, the trace at the end of the 160-ms delay, scaled to each delay’s trace size, was added to the cue-0 state at the end of each of the seven delays. The push of a response was its change in the readout’s margin of the cued over the other target’s move. Bits were the mutual information between the cue and the first push (toward the cued target, the other target or neither) over the trained timings, and spikes per trial came from the mean firing rate.

For the cue flow by cell type (Figures 2H and 2K; Figure 4I; Figure S4G), we rebuilt the effective connectome of the scene-1 readout learner with the simulator’s rules and checked it against its record by hash; the learned group gains were not applied. The cue drive from one cell type to another at the first decision was the sum, over their connections, of the effective weight times the presynaptic difference in spikes between the cues.

#### Ensembles, populations and connection rules

The cue ensemble of a scene was the 1% of cells with the largest absolute cue difference in voltage at the first decision, after the 160-ms delay and 4 input windows. Its 20 matched sets each drew as many cells outside the ensemble and the input cells, with the ensemble’s counts in every stratum of cell type, synaptic distance from the input, degree quintile and readout role; coarser strata stood in where a stratum ran short. Matched synapse sets drew as many connections outside the ensemble, by the strata of both partners and the sign. Silencing gated the output of a set during the first decision; synapse removal set its internal connections to zero for the whole trial. Each ensemble, found in scene 1 or scene 2, was tested in both scenes at the trained timings, against its matched sets.

Population removal deleted half or all of the Kenyon cells (4,064) or of the GABAergic antennal-lobe local neurons (147). Each removal had 10 null sets of the same size, from cells that were neither inputs, readouts nor targets, matched by synaptic distance from the input and degree quintile. It was evaluated at the trained timings in five memory models: the four learners and a scene-1 readout trained with the target hidden or shown (126 trained histories in all).

The connection-rule variants each changed one rule and rebuilt the effective weights: signs from the presynaptic transmitter alone; a same-type gain of 0.05 or 0.2, or none; or a cap of 30 or 120 synapses, or none. The default rules, rebuilt the same way, reproduced the original weights bit for bit. Under each of the 8 rule sets, three replicates of the scene-1 readout learner were trained, and the tests of healthy circuits (below) were repeated.

#### Routes, kicks and excitability

Routes were directed walks of two or three synapses from the visual input cells to the readout cells in the stored connectome, with signs ignored; a cell’s traffic was the number of routes through it. Random removal deleted 30%, 60% or 90% of the cells that were neither inputs nor readouts, in seeded draws, and hub-first removal deleted them in order of degree. A kick added one spike to one cell at cue offset, by raising its voltage above threshold for one step. The kicked cells were five drawn from those that were neither inputs nor readouts, at two trained timings and for both cues (20 kicks).

Threshold sweeps shifted the threshold of every cell by the same amount and measured the mean firing rate under the input of the memory task. Training across threshold states drew, for each training episode, one threshold offset for all cells from a Gaussian with a standard deviation of 1 μV, in 5 replicates of the scene-1 readout learner. Such a run had learned only when it solved every trained history without an offset and at offsets of minus and plus one standard deviation.

Robustness was compared with the 10 replicates trained in one state by three measures. The access window was the uniform lowering of the threshold at which trained memory use first fell to 0.5 or below (log-linear interpolation). Cell-loss tolerance was the share of randomly removed cells at which it did so, averaged over three draws. The third was memory use at two held-out offsets, at *±*0.674 standard deviations.

#### Joining single-cell atlases

Each model cell took an atlas unit through FlyBrain’s type maps, by tier: exact type name, alias, fuzzy match or cell class, preferring the best tier and then a fixed order of sources. A unit’s expression was the mean of log-normalized counts over its cells (over its samples for the bulk data of Davis et al.) and the share of cells detecting the gene. In the ventral nerve cord, which no source resolves by type, we bridged through hemilineages. We reprocessed its atlas with its authors’ pipeline in Scanpy: ^53,74^ their quality control, 2,246 variable genes shared by both sexes, regression of replicate, counts and mitochondrial fraction per sex, and 45 principal components.

Leiden clustering ^75^ at a resolution of 17.9 gave 240 clusters, twice the published number. The Hungarian algorithm matched one new cluster to each published cluster on standardized marker scores, and the others joined their best match. The published hemilineage labels then gave 21 units, each a single hemilineage or a pair that the atlas does not separate, joined to the hemilineage labels of the connectome’s ventral nerve cord cells.

The bridge had to pass three gates declared before it read any atlas cell: recovered against published cluster sizes, the transmitter of each unit against the electron-microscopy prediction, and the enrichment of its markers. Two earlier declared methods failed the size gate, because single-marker clusters absorbed low-quality cells; the third passed all three (Figure 4H).

The panel held 42 genes with their FlyBase identifiers (Table S4). The adaptation index of a unit was the mean *z* score of *SK*, *slo*, *KCNQ* and *Shab* over the mapped units of its source, and its Kv4 weight was its relative *Shal* expression. Trace carriers were the cells with a cue difference in adaptation at the first decision, after the 160-ms delay and 4 input windows. Their mean *SK* and adaptation index were tested against matched sets, label permutations within strata of superclass and source, and against a mapping null that permuted values across mapped types within the same strata, one-sided with 10,000 draws. The ventral nerve cord test used only the hemilineage units.

Heterogeneous brains scaled the adaptation increment by 0.5 in the mapped cell types of the lowest third of the adaptation index and by 1.5 in the highest third (609 types). A null brain permuted these factors across types. Each brain was learned by 10 replicates of the scene-1 readout learner and 5 of the scene-2 learner.

#### Fly pathologies

A literature search found 24 fly pathologies with a measured change in identified cells (Table S1). Each was graded 0 to 2 on six criteria, C1 to C6 (Figure 7A). The first three asked for evidence in flies, a mechanism measured and mappable to model cells, and a change that the model’s operators can express. The last three asked for healthy circuits that behave like the fly’s, the fly phenotype reproduced at the calibrated dose, and its direction kept at half and one and a half times the dose and in a second trained brain. A case was retained if it passed the first five, ranked by its total, with at most four retained. Ties were broken by reach into the planned assays of learning, memory and movement, then by the planned comparison of Aβ42 with tau.

Operators changed one measured property of identified cells with mechanisms the model already has: a threshold offset, a scaling of the adaptation increment, of presynaptic release or of synaptic input, and cell removal. Each is the identity at zero dose. The tests of healthy circuits, on two brains, were sparse Kenyon-cell odor coding, ^79^ its loss without APL feedback, ^80^ the lateral inhibition of the antennal lobe, ^81^ network stability, the giant fiber’s response to a looming stimulus ^82^ and the absence of any effect of monoaminergic cells.

The Kv4 loss that Aβ42 causes in flies ^46^ was a uniform threshold offset of −3.22 mV on every cell at dose 1. This offset gives the fly’s ratio of the ramp current needed for the first spike (0.565 in the model, 0.54 in the fly). The other retained cases removed 17% of the Kenyon cells, scaled the release of GABAergic antennal-lobe local neurons by 0.5, or raised the Kenyon-cell threshold to match their measured fall in odor response. Each had a size-matched random control.

The lesion after learning added the operator to a trained model. Learning under the lesion trained with the operator on from the start, in 6 replicates per dose of the two scene-1 learners. Relearning applied it to a healthy-trained readout for 100 more updates, and the recovered share was the gain in solved histories over the loss. Late restoration removed it from models that had learned under it. ^47^

The *Shal*-patterned lesion multiplied the offset of each cell by its Kv4 weight, imputed by class for unmapped cells and scaled to a mean of 1. The aging operator converted the change in summed potassium-channel transcripts between young and old flies, ^48^ replicated in the Aging Fly Cell Atlas, ^51^ into threshold offsets through the Kv4 calibration.

The olfactory assay read which of two odors a lesioned brain had seen after a delay, without a trained policy. Olfactory receptor neurons fired as Poisson processes, so trials differed only by input noise, and a shrinkage linear discriminant read the odor in 6-fold cross-validation. The read-out sets were the readout features, the mushroom-body output neurons, the Kenyon cells and the projection neurons, for similar and dissimilar odor pairs at delays up to 640 ms. Cases were compared by the distance between their profiles of decoding accuracy, against a null that swapped the paired trials between cases (10,000 draws). In the Alzheimer’s disease atlas, ^52^ expression was averaged per library and regressed by ordinary least squares on genotype, age and sex.

#### Writing and reading the trace in different brain states

The lesion link used the scene-1 readout learner, whose brain is fixed, so its brain states depend only on the input and the lesion. At the end of the delay of each trained history, the full simulator state was replaced by the state the other brain reached there on the same input, and the trial continued in the second brain. This was done in both directions, between the healthy brain and the lesioned brain at doses 0.5 to 2.

The readouts were those trained healthy (10), those trained under the lesion (6 per dose), and readouts relearned under it (3 per dose), rebuilt by replaying their 100 updates exactly. Before the declaration, the replica reproduced the recorded assessments, and transplants within one brain changed no decision. In the run, every condition that involved no transplant reproduced its earlier record.

#### Renders and videos

The neuron renders projected the soma positions, neuron skeletons and neuropil meshes of the connectome release ^2^ with our own renderer. Charts were drawn with Matplotlib. Videos were encoded with FFmpeg (H.264) with fixed settings, so that a rebuild is identical to the byte. In Video S2 and Video S3 the trace was rebuilt from the recorded spikes with the model’s adaptation rule and checked against the record at every millisecond.

#### Reproducibility

The simulation is deterministic: every input was a constant current or a seeded draw, and reruns were bit-identical on every GPU we used (NVIDIA GeForce RTX 5090), which we checked before each campaign. An independent process reloaded and verified every trained model; Table S3 lists every training run and evaluation with its outcome and record. The designs and test families were declared before their runs, in test plans and hash-chained amendments; families added after results were post hoc and never primary, and results were never overwritten. The analysis reruns from the records to identical outputs, a manifest pins every result, figure and video by its SHA-256, and Data S1 holds the values behind every figure panel.

### Quantification and statistical analysis

The independent unit of a training result was a run, trained from its own replicate seed. Within one model, the unit was a history under fixed parameters, and maps were the unit of the planning ladder. Cells, time points, timings and checkpoints were never treated as replicates. The *n* of each test is given in the figure legends and in Table S2.

Across runs we used exact sign-flip tests of mean differences, exact Mann–Whitney tests and a stratified log-rank test of learning times. Within models we used exact McNemar tests on paired histories and Page trend tests over doses. Correlations were Spearman’s, with permutation tests; matched-set and mapping nulls gave *P* = (*k* + 1)/(*N* + 1) from *N* draws. Claims of no difference used two one-sided tests, with Tango’s test for paired proportions ^83^ and a margin of 5 percentage points.

Intervals for ratios of geometric means came from bootstrap resampling (95%). Resampling enumerated every case when *n* was at most 20 and otherwise used 10,000 draws, each seeded from one analysis seed and the test’s name. *P* values are two-sided unless stated. Each test family was declared before its tests were computed. Within a family, *P* values were adjusted by Holm’s method for primary families and by the Benjamini–Hochberg method for secondary and exploratory ones, always over the complete declared family.

Table S2 lists every declared test the paper reports, with its family and correction. The outcomes below appear only here and in Table S2:

- From a neutral brain, training the readout alone learned the planning maps faster than training the brain and readout together, and all 24 runs learned (stratified log-rank *P* = 6.3 × 10^−5^).
- By the stagnation rule, 9 of 10 runs of the scene-1 brain-and-readout learner, 3 of 7 of the scene-2 learner and 7 of 7 of the scene-3 learner stopped before meeting the learning criterion, each keeping its best model.
- The models trained on single representative planning maps solved none of 8 held-out maps without further updates.
- Replayed from the original start, none of the 650 models saved at curriculum stages solved its map, as the curriculum ensures, and their first decisions showed no trend toward those of the final model.
- Adaptation alone, transplanted at the end of the trained 160-ms delay, did not reproduce the decision (cosine 0.33), and responses to scaled traces changed direction with their size.
- Shifting the state toward the other cue’s changed decisions more than norm-matched scrambled shifts in only 2 of 6 models (Holm).
- Swaps disrupted histories with the target hidden and shown alike (*P* = 0.38), against the predicted difference.
- In earlier follow-up analyses, state swaps were specific to the cue but not to memory, and most claims of absence were inconclusive.
- No linearization of the network was validated, so no spectral measure was interpreted.
- Removing as many matched random cells as the populations cost 23 to 36 percentage points more of the trained histories; removing the descending neurons, the positive control, left 57 of 126 solved and silenced the giant fiber.
- At a same-type gain of 0.05, the strict lock was absent (untrained use +0.19 to +0.35), and without same-type damping the mushroom-body tests of healthy circuits failed.
- With mapping by cell type alone, the carriers did not exceed matched sets, an effect of source composition, and the adaptation index did not exceed the mapping null (*P* = 0.068 to 0.14).
- Heterogeneous brains did not differ from their null brains (adjusted *P* = 0.091 to 0.14); all 5 scene-2 runs were positive, at adjusted *P* = 0.057.
- Jittered training lowered trained-timing use by 0.23 and took 1,090 more updates. One new scene generalized partly to untrained timings, and for the scene-1 brain-and-readout learner the test of untrained use gave Holm *P* = 0.081.
- A uniform lowering of the threshold by 0.17 to 0.25 μV halved memory use (per replicate), while the mean rate stayed at 0.35 to 0.37 Hz and the network ran away only from 6.5 mV.
- Under such shifts the spike trajectories diverged only once input resumed, and the share of trained histories solved fell from 0.92 to 0.84 and 0.63 at lowerings of 10^−4^ and 3 × 10^−4^ mV.
- A persistent lowering of 0.3 μV left the delay silent; from 11 ms after input resumed the network diverged, and 4,659 cells had diverged by the end of the first decision, motor and efferent cells most often.
- The dose series did not identify the dose of the Kv4 lesion that halves memory, which lay below a lowering of 0.01 mV.
- Random removal of 0.13% of the cells halved behavior (four-parameter logistic fit).
- Training across threshold states reached trained use of 0.89 and untrained use of 0.11, with 3.6 times the updates.
- In the olfactory assay, targeted lesions did not exceed random ones. Relearning showed no dose trend, relearning with the brain recovered less (−0.29), and no per-dose test of learning under the lesion was significant after correction (smallest adjusted *P* = 0.19).
- At doses 0.5 and 1, the *Shal*-patterned and uniform lesions were not shown to be equivalent.
- In the Aging Fly Cell Atlas, the change in Kenyon-cell potassium-channel transcripts had the opposite sign to that in the primary atlas.
- Reading in the state of learning beat writing in it for readouts trained under the lesion (Holm *P* = 3.0 × 10^−4^), but not for readouts trained healthy (1.00) or relearned (0.26).

## Notes

https://doi.org/10.5281/zenodo.23629827

https://doi.org/10.5281/zenodo.23629835

https://github.com/Hanziwww/fly-connectome-silent-memory

### Competing Interest Statement

The authors have declared no competing interest.

