## Supplementary material for "Brain state locks silent memories in a trained fly connectome": Document S1 - Figures S1-S7 and Table S1

### Figure S1. The model, what training changes, and its validation, related to Figure 1

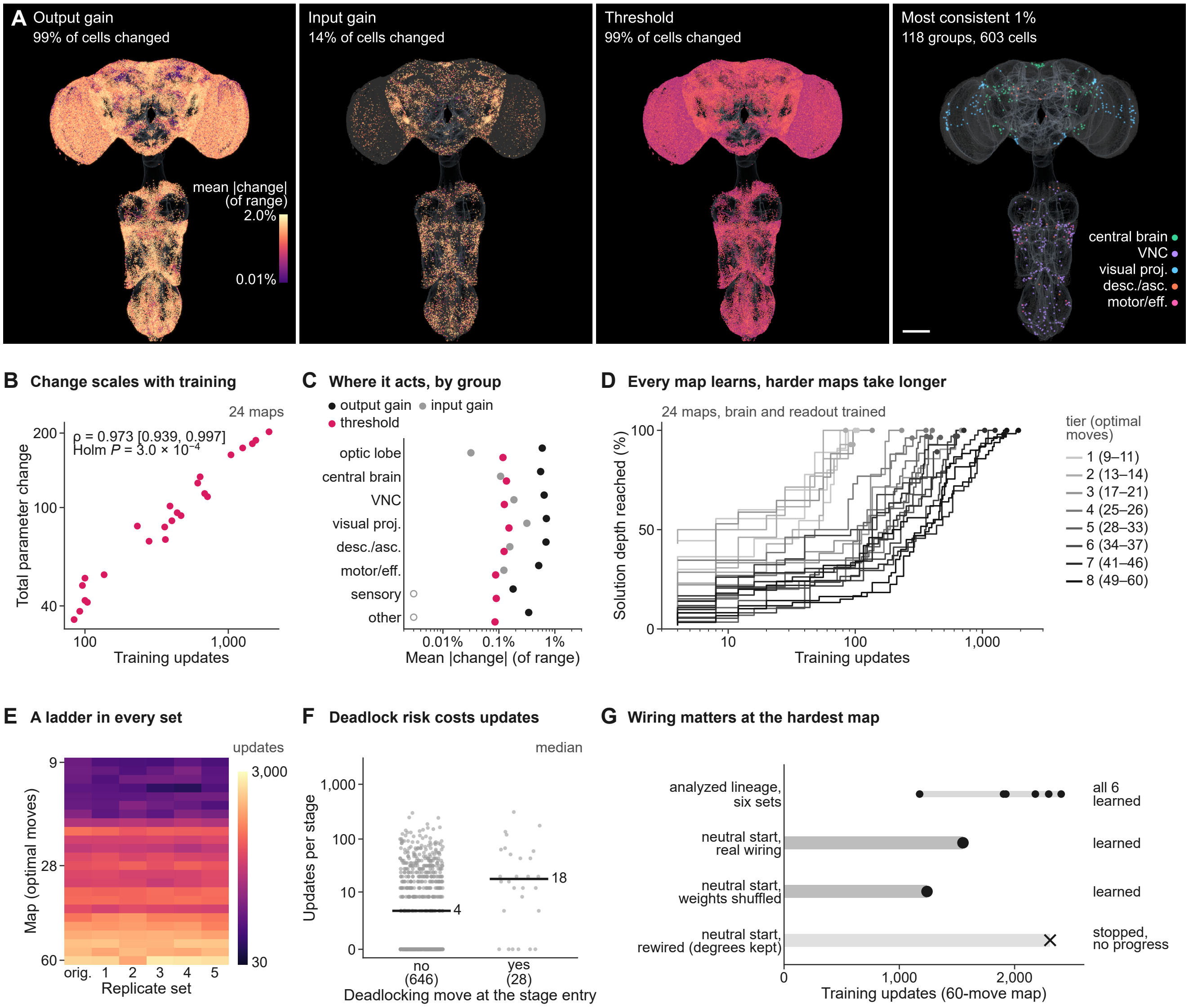


#### H The model at a glance

| Network | Neurons | Trained |
|---|---|---|
| **167,106** neurons (all of the CNS) | **−52 / −45 mV** rest / threshold | **35,310** brain parameters |
| **25.58 M** connections (wiring fixed) | **20 / 5 ms** membrane / synaptic τ | **3 × 11,770** gains in, out, threshold |
| **14.78 M / 9.88 M** excitatory / inhibitory | **0.275 mV** per synapse (cap 60) | **2,052** readout weights and biases |
| **11,770** cell-type groups | **+1.5 mV, 200 ms** adaptation per spike, τ | **512** readout features |
| **3,200** visual input cells | **2.2 / 1.8 ms** refractory / delay | **4** actions |
| **4,096** readout cells (256 × 16) | **1 ms** time step | **40 ms** decision window |

#### I Bit-exact on every GPU

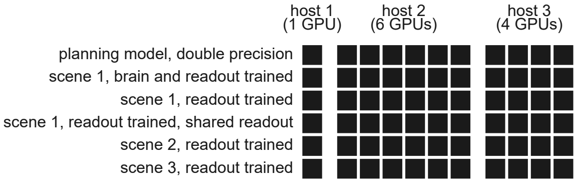

**Figure S1. The model, what training changes, and its validation, related to Figure 1**

(A) Where training acts in the planning model: each cell by the mean absolute change of its cell-type group's output gain, input gain and threshold over the 24 maps (log color), and the cells of the most consistently changed groups. Scale bar, 100 μm. (B) Total parameter change against training length, one dot per map; Spearman ρ with a BCa bootstrap interval, permutation test, Holm-adjusted. (C) Mean absolute change by cell group and parameter. (D) Every map's learning curve: solution depth reached against training updates, by difficulty tier. (E) Updates to solve each map (rows by optimal moves) in the original set and five replicate sets. (F) Updates per stage for stages entered with or without a deadlocking move available; bars, medians. (G) The hardest map: training updates of the analyzed lineage in six sets, and from a neutral start with the real wiring, with its weights shuffled and rewired with degrees kept (one run each); cross, stopped without progress. (H) The model's parameters at a glance. (I) Bit-exact replay of every analyzed model on every graphics processor of the three training hosts.

#### Figure S2. The silent trace in every learner: written, held, read, related to Figure 2

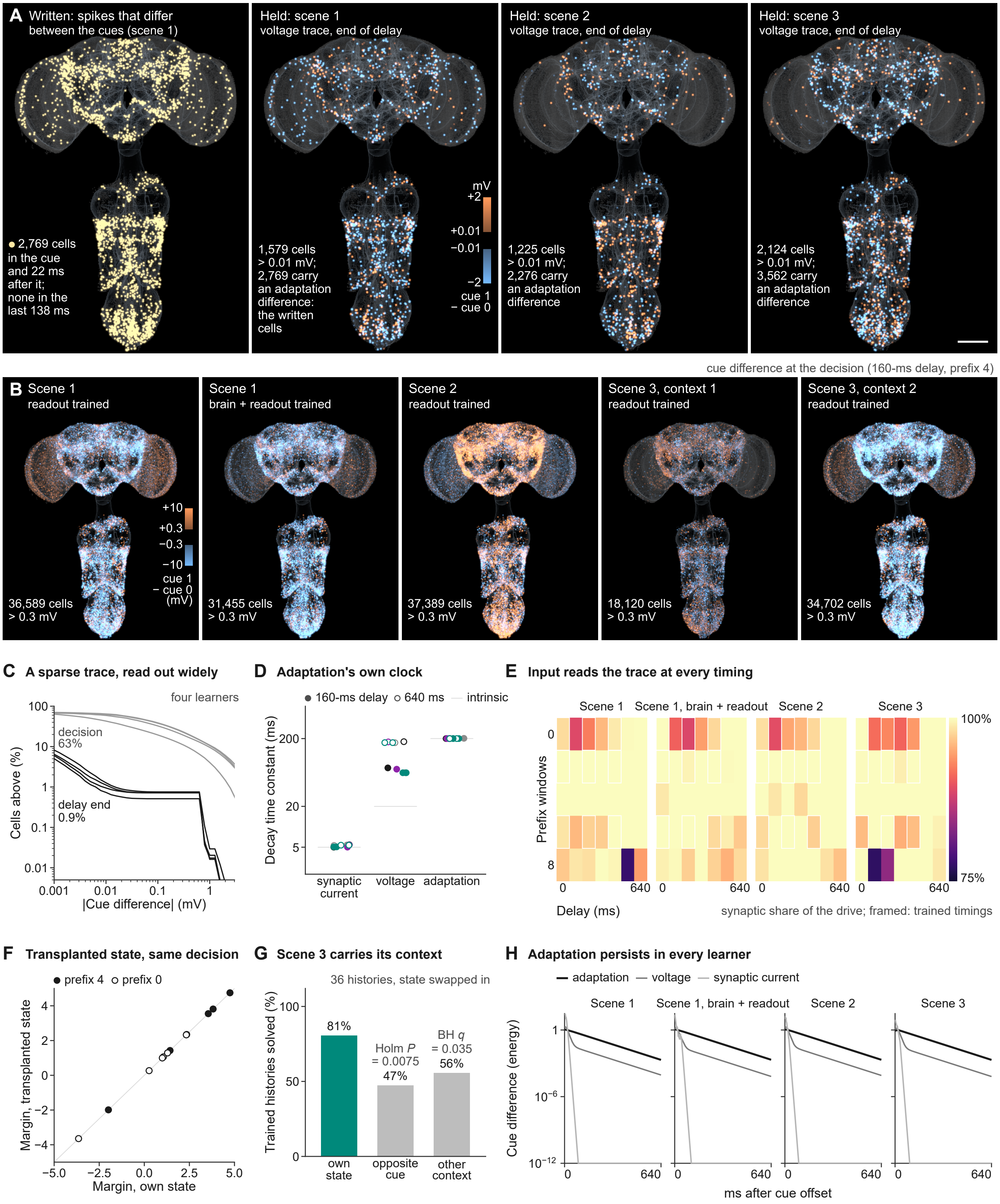

**Figure S2. The silent trace in every learner: written, held, read, related to Figure 2**

(A) Where the cue writes the trace: the cells that spike differently during the cue (scene 1), and the voltage difference held at the end of the delay in scenes 1–3, with the cells that carry an adaptation difference. Scale bar, 100 μm. (B) The cue difference at the decision in five state sets: scene 1 with the readout or the brain and readout trained, scene 2, and scene 3 in each of its two contexts. (C) The share of cells above each absolute cue difference, at the end of the delay and at the decision, four learners. (D) Time constants of the cue difference by state variable, in every learner, timing and context; line, the intrinsic time constant. (E) The synaptic share of the decision-time drive on the grid of delay and prefix input, four learners; framed, trained timings. (F) The decision margin with the transplanted state against the native margin, at every delay, without and with prefix input. (G) Scene 3 holds its context with the cue: trained histories solved with the own state, the opposite cue's state or the other context's state; exact tests, Holm- and BH-adjusted. (H) The cue difference by state variable through the 640-ms delay, in every learner.

**Figure S3. No circuit is special, in every scene and under every connection rule, related to Figure 3**

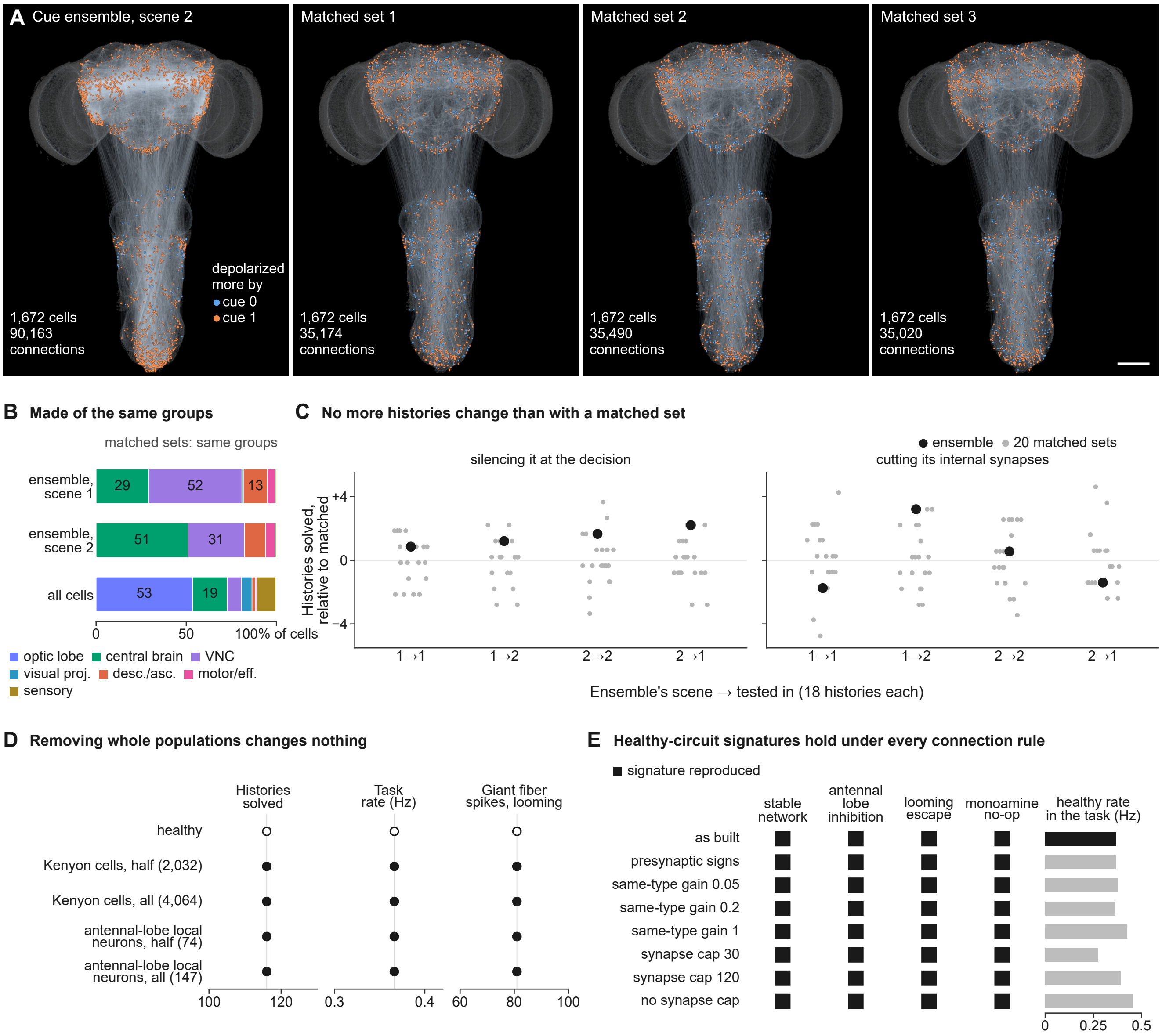


(A) The scene-2 cue ensemble (the 1% of cells with the largest cue difference at the decision) and its internal connections, beside three of its 20 matched sets drawn by the same rule. Scale bar, 100 μm. (B) The cell groups of the scene-1 and scene-2 ensembles, which their matched sets share by design, against all cells. (C) Trained histories solved after silencing the ensemble or cutting its internal synapses, relative to the mean of the matched sets (gray), by the ensemble's scene and the scene tested. (D) Removing whole populations: trained histories solved, the network's rate and the giant fiber's looming response, against healthy. (E) The signatures of healthy fly circuits used by the disease study (a stable network, antennal-lobe inhibition, looming escape and the monoamine control) and the task rate of the healthy network under every connection rule.

#### Figure S4. The molecular atlas behind the *SK* signature, related to Figure 4

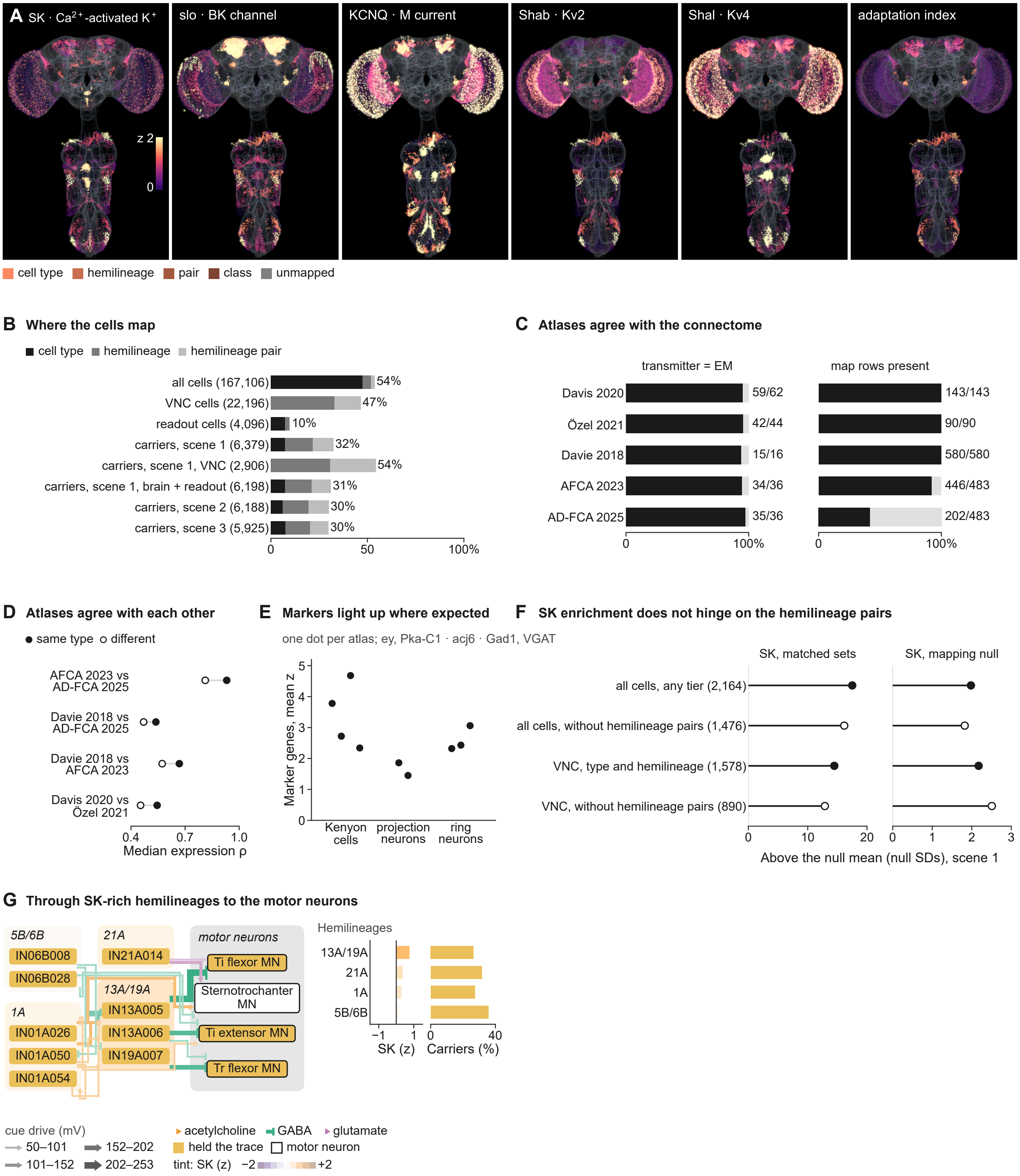

**Figure S4. The molecular atlas behind the *SK* signature, related to Figure 4**

(A) Single-gene in silico stains on the unfolded CNS (*SK*, *slo*, *KCNQ*, *Shab*, *Shal* and the adaptation index): each mapped cell by its atlas unit's z score, with the mapping tier as brightness. (B) The share of each cell set mapped by cell type, by hemilineage and by hemilineage pair. (C) Agreement with the connectome, per atlas: the transmitters of the mapped types against the predictions from electron microscopy, and the map rows present. (D) Agreement between atlases: the expression correlation of units mapped to the same cell type, against different types. (E) Marker genes in the mapped units of Kenyon cells, projection neurons and ring neurons, one dot per atlas. (F) *SK* enrichment of the scene-1 carriers with and without the hemilineage-pair tier, against matched sets and the mapping null. (G) The cue's routes to the motor neurons by hemilineage: modules tinted by atlas *SK* (z); gold, types that held the trace; outlined, motor neurons; width, the cue drive at the first decision; beside it, each hemilineage's *SK* and carrier share.

#### Figure S5. Access locked to the timing of learning, at every delay, related to Figure 5

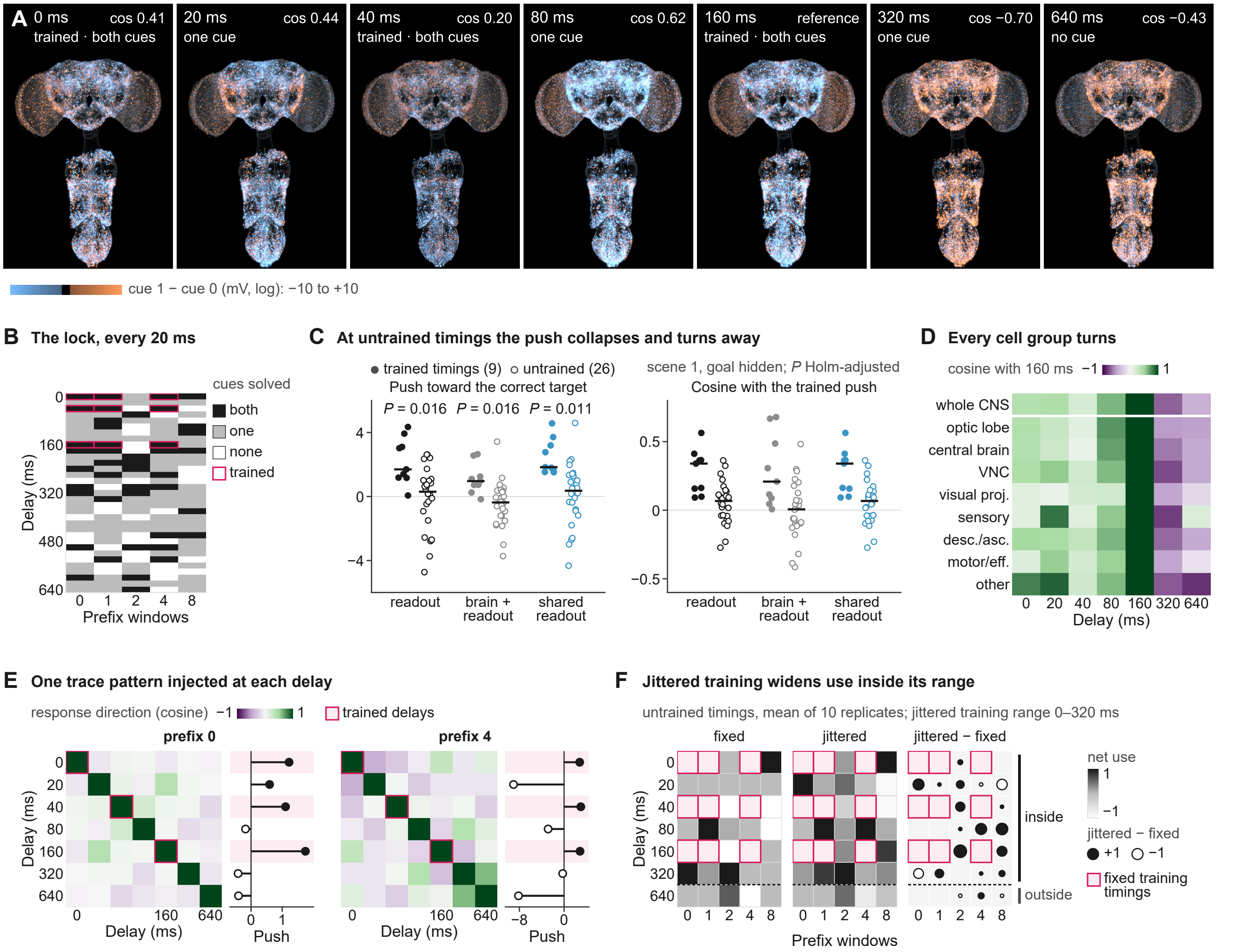


(A) The decision-time cue difference at all seven delays (four prefix windows; scene 1, readout trained), each with its outcome and its cosine with the trained 160-ms pattern. (B) Cues solved at every delay of the dense scan and every prefix input; framed, trained timings. (C) At untrained timings the push toward the correct target collapses and its direction turns away from the trained push, in the three readouts of scene 1 in which the collapse survives correction; bars, medians; Holm-adjusted *P*. (D) The cosine with the 160-ms pattern within each cell group, by delay. (E) One trace pattern injected at seven delays, without and with prefix input: the cosine between response directions, and each response's push; framed, trained delays. (F) Untrained use after training at fixed or jittered timings, timing by timing (means of 10 replicates per arm), and their difference.

#### Figure S6. Many routes, fading kicks and a wider window, in detail, related to Figure 6

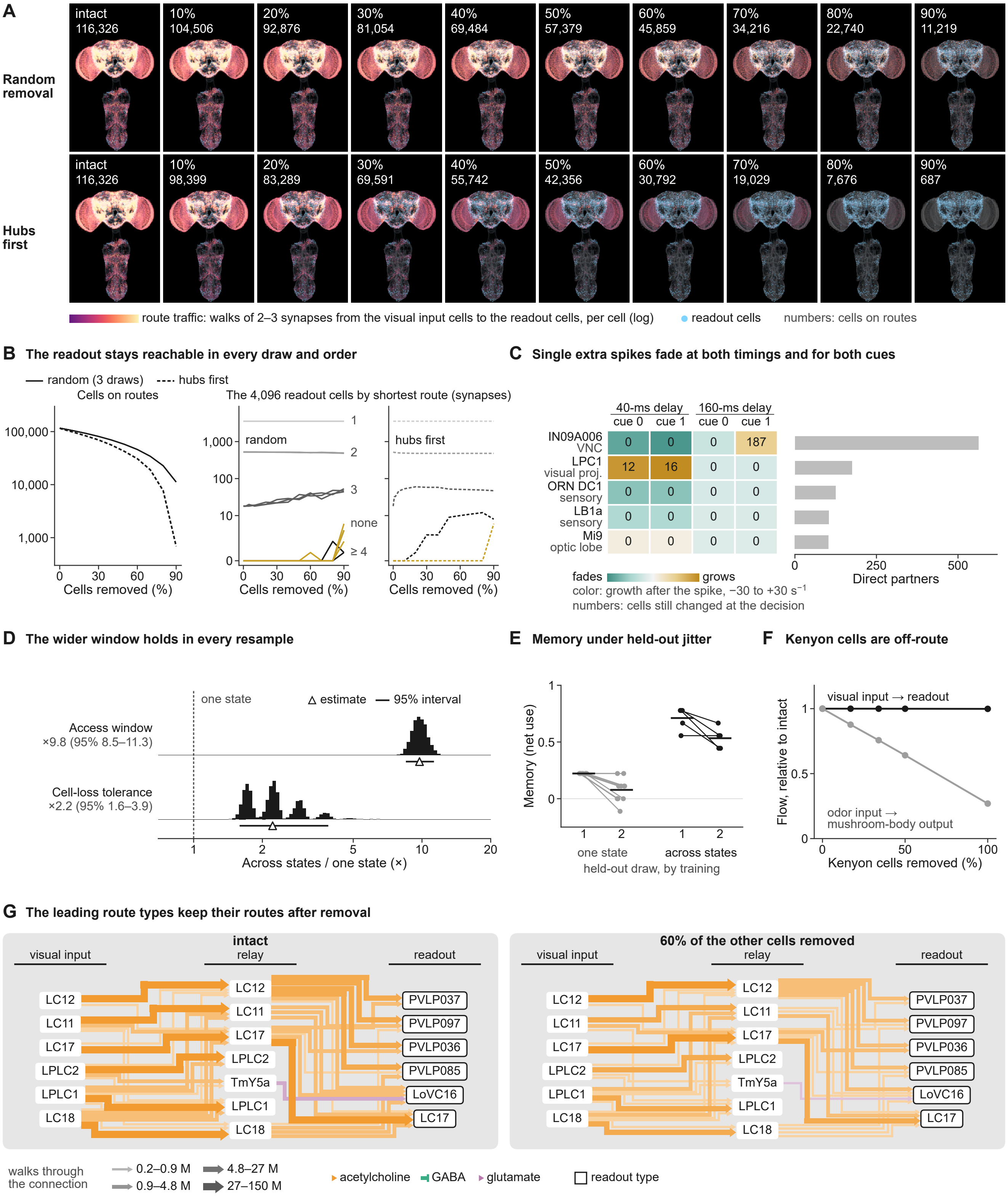

**Figure S6. Many routes, fading kicks and a wider window, in detail, related to Figure 6**

(A) The route network at every tenth of removal, random (the first of three seeded draws) and hubs first (cells in descending order of connections); numbers, cells on routes. (B) Cells on routes, and the readout cells by shortest route, in every draw and both orders. (C) Single extra spikes: per kicked cell, the growth of the difference after the spike and the cells still changed at the decision, at two delays and for both cues, with the cell's direct partners. (D) Bootstrap resamples of the access window and the cell-loss tolerance, across states over one state; triangles, estimates; lines, 95% intervals. (E) Memory under each held-out jitter draw, replicate by replicate; bars, means. (F) Flow capacity from the visual input to the readout and from the odor input to the mushroom-body output, with Kenyon cells removed. (G) Typed routes from the visual input to the readout, intact and with 60% of the other cells removed: the relay types with the most route traffic, the input and readout types that use them most, and every connection among them above a threshold; width, walks of two or three synapses through the connection; color and arrowhead, the sender's transmitter.

#### Figure S7. Disease steals access, not memories, in detail, related to Figure 7

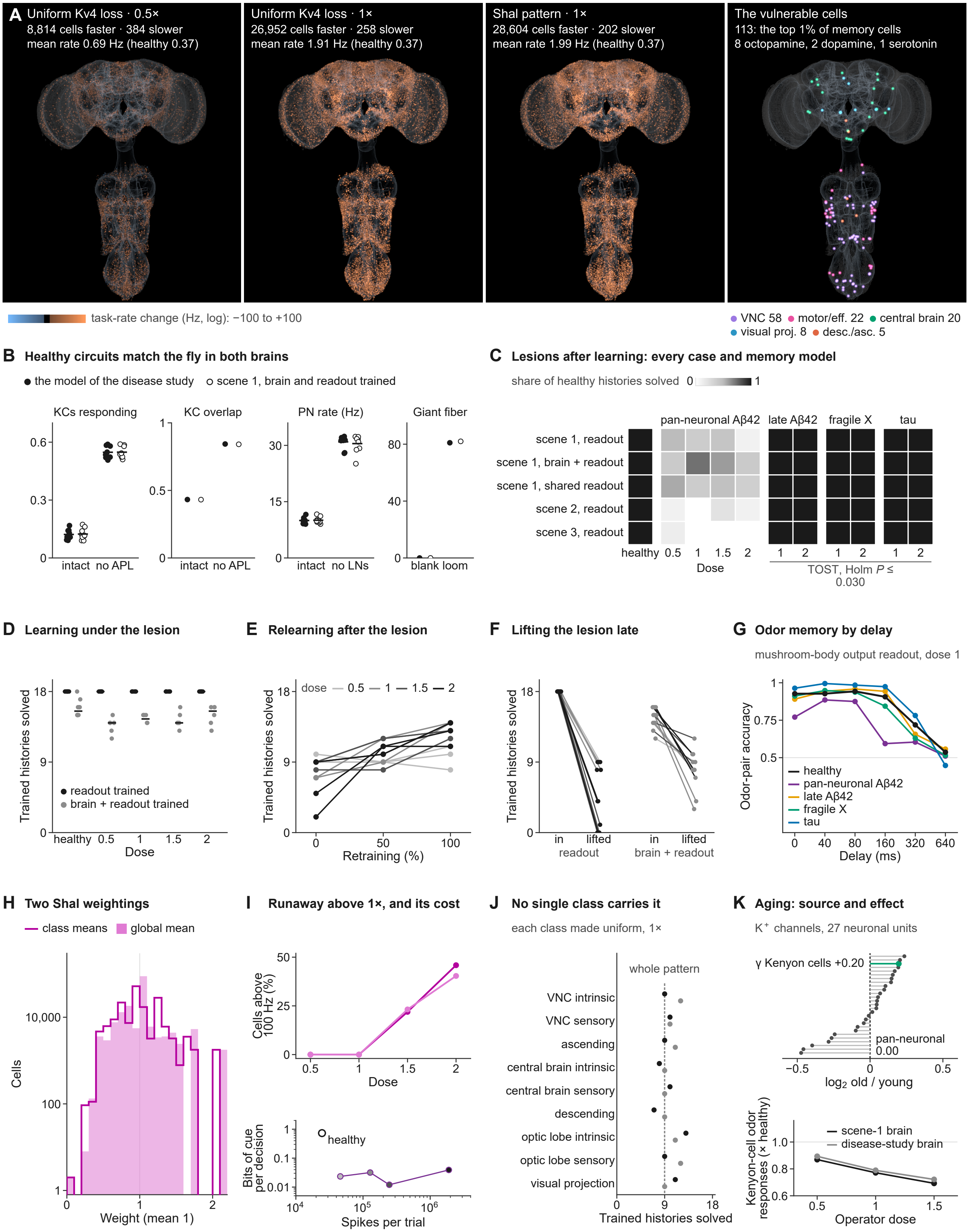

**Figure S7. Disease steals access, not memories, in detail, related to Figure 7**

(A) Each cell's change in task rate under the uniform Kv4-loss lesion at half and full dose and under the *Shal* pattern of the same mean, and the vulnerable cells (the 1% of memory-relevant cells with the largest change) by group. (B) The healthy circuits match the fly in both brains: sparse Kenyon-cell coding broadened by APL removal, projection-neuron rates raised by removing the local neurons, and the giant fiber's looming response. (C) Lesions after learning in every retained case and memory model; equivalence tests (TOST), Holm-adjusted. (D) Learning under the lesion, run by run; bars, means. (E) Relearning after the lesion, replicate by replicate, with the readout retrained. (F) Lifting the lesion late removes access in every run. (G) Odor memory by delay in every case, read from the mushroom-body output. (H) The two *Shal* weightings of one mean loss: class means and the global mean. (I) Above full dose both weightings run away (cells above 100 Hz); below, spikes per trial against bits of cue per decision. (J) Trained histories solved when each superclass's *Shal* weighting is made uniform in turn (descriptive). (K) The aging operator: its source, the summed potassium-channel expression of old over young flies in neuronal atlas units,[1] and its effect on Kenyon-cell odor responses in two brains.

**Table S1. The credibility screen of fly pathologies, related to Figure 7**

Each candidate pathology of the fly, with the change measured in flies and the operator that expresses it in the model. C1–C3 grade, from 0 to 2, the evidence in flies, a mechanism measured and mappable to model cells, and a change that the model's operators can express; candidates that passed all three entered the tests in the model. C4, the healthy circuits behave like the fly's (one set of tests in two brains); C5, the fly phenotype is reproduced at the calibrated dose (graded); C6, its direction holds at half and one and a half times the dose and in a second trained brain. Total, the sum of the six grades (a pass of C4 or C6 counts 2), which ranked the cases that passed C1–C5; at most four were retained. Dashes, not tested. Sources are given by DOI.

| Fly condition | Change measured in flies | Operator in the model | C1 | C2 | C3 | C4 | C5 | C6 | Total | Outcome | Sources (DOI) |
|---|---|---|---|---|---|---|---|---|---|---|---|
| Fragile X (*dfmr1* null) | Weaker GABAergic lateral inhibition in the antennal lobe; broader odor tuning | Release of GABAergic antennal-lobe local neurons scaled down | 2 | 2 | 2 | pass | 2 | pass | 12 | Retained | 10.1016/j.cub.2017.02.065<br>10.1016/j.nbd.2014.01.008 |
| Aβ42, late (25 to 50 days) | Kenyon cells lost first, then other neurons | Kenyon cells removed | 2 | 2 | 2 | pass | 1 | pass | 11 | Retained | 10.1371/journal.pgen.1005025<br>10.1073/pnas.0400895101<br>10.1371/journal.pone.0001703<br>10.1016/j.celrep.2018.11.098<br>10.1186/s40478-016-0333-4 |
| Aβ42, pan-neuronal | Kv4 (A-type $K^+$) channel degraded; less current needed for the first spike | Threshold lowered in every cell | 1 | 1 | 2 | pass | 2 | pass | 10 | Retained; the only case that reaches the visual tasks | 10.1371/journal.pgen.1005025<br>10.1016/j.neuroscience.2017.06.008<br>10.1016/j.cub.2015.01.016<br>10.3389/fncel.2019.00260 |
| Human tau or Aβ42 in Kenyon cells | Depolarization-evoked $Ca^{2+}$ influx down by about half | Threshold raised in Kenyon cells | 1 | 2 | 2 | pass | 1 | pass | 10 | Retained for the planned comparison of Aβ42 with tau | 10.3389/fncel.2019.00409<br>10.3389/fncel.2019.00260 |
| Aβ42 in mushroom-body neurons | Kv4 current lost in Kenyon cells | Threshold lowered in Kenyon cells | 1 | 2 | 2 | pass | 1 | pass | 10 | Passed; not retained (rank) | 10.1371/journal.pgen.1005025<br>10.1016/j.neuroscience.2017.06.008 |
| Aβ42, early (0 to 2 days) | More spontaneous transmission onto projection neurons | Release onto projection neurons scaled up | 1 | 2 | 2 | pass | 1 | pass | 10 | Passed; not retained (rank) | 10.1016/j.celrep.2018.06.029<br>10.1007/s12264-012-1267-x |
| Aβ42, 9 to 13 days | Less spontaneous transmission onto projection neurons | Release onto projection neurons scaled down | 1 | 2 | 2 | pass | 1 | pass | 10 | Passed; not retained (rank) | 10.1016/j.celrep.2018.06.029<br>10.1016/j.nbd.2012.11.006<br>10.1016/j.cub.2015.01.016 |
| Normal aging (3 vs 30 days) | Larger active zones and more release at projection neuron to Kenyon cell synapses | Release raised at these synapses | 2 | 2 | 2 | pass | 0 | fail | 8 | Excluded (C5): Kenyon-cell responses doubled; in flies they stay flat or fall | 10.1371/journal.pbio.1002563<br>10.1038/nn.3512<br>10.1371/journal.pbio.3001730<br>10.1002/cne.23798<br>10.1038/s41467-019-09262-2 |
| Human tau in output neurons M4 and M6 | Depolarization-evoked $Ca^{2+}$ influx up about 2.7-fold | Threshold lowered in these output neurons | 1 | 2 | 2 | pass | 0 | fail | 7 | Excluded (C5): the fly's increase lies beyond the operator's range | 10.3389/fncel.2019.00409 |
| Normal aging (1 to 5 vs 25 to 30 days) | *KCNQ* (M-current) transcripts fall in the brain | Threshold lowered in α/β Kenyon cells | 1 | 1 | 2 | pass | 0 | fail | 6 | Excluded (C5): no change at the circuit level, and no matched assay | 10.1371/journal.pone.0062445<br>10.1016/j.cub.2015.01.016 |
| Aβ42 inside identified neurons | Lower release probability, fewer $Ca^{2+}$-channel clusters, less Bruchpilot | Release scaled down | 1 | 1 | 2 | – | – | – | – | Deferred (C3): no published magnitude to calibrate | 10.1016/j.nbd.2012.11.006<br>10.1523/JNEUROSCI.3699-09.2010<br>10.1016/j.celrep.2018.06.029 |

| Fly condition | Change measured in flies | Operator in the model | C1 | C2 | C3 | C4 | C5 | C6 | Total | Outcome | Sources (DOI) |
|---|---|---|---|---|---|---|---|---|---|---|---|
| Human tau isoforms, giant fiber system | More response failures at high frequency; slower conduction | Stronger short-term depression at the identified synapse | 1 | 1 | 2 | – | – | – | – | Deferred (C3): its magnitude needs the full text | 10.1016/j.nbd.2018.12.004 10.1002/dneu.20388 10.1523/JNEUROSCI.3699-09.2010 |
| Aβ42 inside a flight motor neuron | Faster synaptic fatigue at high frequency | Stronger short-term depression; release scaled down | 1 | 1 | 2 | – | – | – | – | Deferred (C2, C3): weak mapping, no magnitude | 10.1523/JNEUROSCI.3699-09.2010 10.1016/j.nbd.2012.11.006 |
| Tau in Kenyon cells, several isoforms | Less depolarization-evoked $Ca^{2+}$ influx; learning impaired | Threshold raised in Kenyon cells | 1 | 1 | 2 | – | – | – | – | Merged with the tau case in Kenyon cells (the same measurement) | 10.1101/lm.70804 10.3389/fncel.2019.00409 10.3389/fncel.2019.00260 10.1523/JNEUROSCI.1374-22.2023 10.1523/JNEUROSCI.1490-09.2010 |
| Normal aging (2 vs 60 days) | Dopaminergic neurons preserved | No change | 2 | 2 | 2 | – | – | – | – | Control: no change to reproduce | 10.3389/fnins.2010.00205 10.1093/sleep/32.8.984 10.1002/gene.20106 |
| Aβ42, tau or α-synuclein in photoreceptors | Photoreceptors lost (Aβ42) | Photoreceptors removed | 1 | 2 | 2 | – | – | – | – | Not tested: confined to the retina, which the visual task input bypasses | 10.1186/s40478-016-0333-4 10.1371/journal.pone.0001703 |
| Normal aging | Mushroom-body fiber number declines after 3 to 4 weeks | Kenyon cells removed | 1 | 1 | 2 | – | – | – | – | Excluded (C1): a single early study, contradicted since | 10.3109/01677068409107077 10.1371/journal.pbio.1002563 |
| Human tau, adult-onset, pan-neuronal | Slower learning; protein-synthesis-dependent memory impaired, reversibly | None available | 2 | 0 | 0 | – | – | – | – | Excluded (C2, C3) | 10.1523/JNEUROSCI.1374-22.2023 10.1016/j.nbd.2018.12.004 |
| α-Synuclein with sleep deprivation | Persistent loss of short-term memory that requires dopamine receptors | None available | 1 | 0 | 0 | – | – | – | – | Excluded (C2, C3) | 10.1093/sleep/32.8.984 |
| Normal aging (10 vs 30 days) | Fewer contacts from DPM neurons onto α/β Kenyon cells | None: DPM output carries no weight in the model | 2 | 1 | 0 | – | – | – | – | Excluded (C3) | 10.1073/pnas.1118126109 10.1523/JNEUROSCI.0978-14.2015 |
| α-Synuclein, parkin or PINK1 | Dopaminergic neurons and their synapses lost | None: dopaminergic synapses carry no weight in the model | 1 | 1 | 0 | – | – | – | – | Excluded (C3) | 10.1016/j.celrep.2013.10.032 10.1073/pnas.0501078102 10.1002/gene.20106 10.1093/braincomms/fcab049 10.7554/eLife.105386 10.1021/acschemneuro.2c00277 10.3389/fnins.2010.00205 |
| Fragile X (*dfmr1* null), 7 to 9 days | Less 5-HT2A receptor in the mushroom body | None: no serotonin receptor signaling in the model | 2 | 1 | 0 | – | – | – | – | Excluded (C3) | 10.1038/s41598-025-34492-4 |
| Sleep loss with Aβ-Arctic | Large ventral lateral clock neurons fire more | Threshold lowered in these clock neurons | 1 | 2 | 1 | – | – | – | – | Excluded (C3): the change cannot propagate in the model | 10.1016/j.cub.2015.01.016 10.1016/j.nbd.2019.104507 |
| Human tau in dopaminergic neurons | Vesicular dopamine release lost | None: dopamine release carries no weight | 1 | 1 | 0 | – | – | – | – | Excluded (C3) | 10.1007/s00401-013-1105-x 10.1021/acschemneuro.2c00277 |
