## Supplementary material for "Brain state locks silent memories in a trained fly connectome": Data S1 - Figure source data: README.txt

Data S1. Source data for Figures 1–7 and S1–S7

The values behind every panel of the paper's figures, one table per figure (.csv, UTF-8). Each row belongs to one
panel (column "panel") and one plotted series (column "series"); the other columns hold the values that series
plots and are empty where they do not apply. Several values in one cell are separated by semicolons.

Figure_1.csv     Figure 1. A whole-CNS connectome learns to plan and to remember
Figure_2.csv     Figure 2. Working memory is held in an activity-silent intrinsic trace
Figure_3.csv     Figure 3. The trace is distributed, and no circuit is special
Figure_4.csv     Figure 4. Trace-carrying cells have a molecular signature: SK
Figure_5.csv     Figure 5. Access to the memory is locked to the timing of learning
Figure_6.csv     Figure 6. The wiring is robust; access follows the trained states
Figure_7.csv     Figure 7. Disease steals access, not memories
Figure_S1.csv    Figure S1. The model, what training changes, and its validation
Figure_S2.csv    Figure S2. The silent trace in every learner: written, held, read
Figure_S3.csv    Figure S3. No circuit is special, in every scene and under every connection rule
Figure_S4.csv    Figure S4. The molecular atlas behind the SK signature
Figure_S5.csv    Figure S5. Access locked to the timing of learning, at every delay
Figure_S6.csv    Figure S6. Many routes, fading kicks and a wider window, in detail
Figure_S7.csv    Figure S7. Disease steals access, not memories, in detail
